# Neural plate morphogenesis and pre-patterning enable specification of intermediate progenitors in the spinal cord

**DOI:** 10.1101/2025.01.09.632276

**Authors:** Sandy Nandagopal, Anna Cha, Bill Z. Jia, Hongyu Liao, Caroline Comenho, Galit Lahav, Daniel E. Wagner, Tony Y-C Tsai, Sean G. Tsung-Megason

## Abstract

Spinal cord organization begins with the specification of a series of neural progenitor domains along the dorsal-ventral axis of the neural tube. Patterning at the dorsal and ventral ends is established by canonical morphogen gradients. Intermediate progenitor domains are located distally to morphogen sources yet display similar organization. How this is achieved is unclear and the relevant positional cues have not been identified. Here we show that intermediate progenitor domains are established through a distinct mechanism that operates in the preceding neural plate stage and leverages its multiple signaling inputs and morphogenesis. An integrated spatial atlas of transcription factor expression, signaling activity, and cell movement reveals that the neural plate comprises an array of cell states, defined by combinatorial signaling and morphogenesis-driven transitions between local signaling environments. Specifically, we find that p0 and p1 intermediate progenitors, located immediately outside the morphogen-patterned region, arise from specific precursor states that are established in the posterior neural plate by tailbud-derived Wnt signaling. Convergent-extension movements subsequently drive progenitor specification by displacing precursors out of Wnt/FGF signaling fields while shaping them into stripe-like domains. Thus, by controlling signaling exposure, tissue dynamics convert positional information encoded by signaling history into downstream fate specification. This mode of patterning may operate broadly across developmental contexts, with implications for tissue engineering and patterning-related disorders.

## Introduction

The development of functional tissues and organs relies on the proper specification and organization of diverse cell types. Developing tissues vary widely in shape, size and other features, ranging from a few immotile cells that behave in a highly stereotyped manner^1,2^ to hundreds of motile cells that vary in their individual behaviors between embryos but nevertheless organize robustly in aggregate^3–6^. A central goal in developmental biology and tissue engineering is to uncover the specific mechanisms and strategies that have evolved to pattern diverse tissue contexts.

Morphogen signaling has emerged as the central mechanism for tissue-scale patterning across organs and organisms. Previous work in several systems has shown that long-range spatial gradients of signaling molecules (morphogens) can encode positional information within tissues, which can be interpreted by cells to activate spatially appropriate fate programs. Morphogen-driven mechanisms can direct precise patterning, especially in combination with extracellular regulatory interactions that tune morphogen distribution^7–9^ and intracellular gene regulatory networks that decode various features of signaling including concentration and temporal dynamics ^10–13^.

Yet there are limits to morphogen-mediated patterning. Work in several developmental contexts suggests that individual morphogen systems typically establish only a limited number of distinct spatial domains^10,14,15^. This can at least partly be attributed to the challenge of establishing precise patterns at increasing distance from the source, where the gradient becomes shallower and the relative level of biochemical noise increases, leading to increased positional uncertainty^16,17^. Another challenge is that tissues often undergo growth and morphogenesis concomitantly with patterning. These changes in tissue size or shape can increase noise and temporal variability in the positional information provided by a static morphogen gradient^18,19^. Though compensatory mechanisms for these issues have been identified in some cases^18–20^, significant gaps remain in our ability to explain patterning in even the most canonical developmental systems.

Dorsal-ventral (D-V) organization of the spinal cord exemplifies both the power and limits of morphogen-based patterning. The spinal cord develops from the posterior neural tube, which is patterned along the D-V axis into 11 neural progenitor domains that produce functionally distinct cell types in the mature spinal cord^21^. Previous work has established that Shh secreted from the notochord and floor plate (at the ventral end of the tissue) acts as a morphogen to pattern a broader region in the ventral neural tube^11,14,22^. This is in fact one of the best understood examples of morphogen-mediated patterning. Cells at different distances from the source of Shh experience different temporal profiles of signaling, and decode this information through a gene regulatory network to determine appropriate neural progenitor identities ^11,12,14^. Moreover, at the tissue level, this mechanism can produce the stripe-like patterns observed in the tissue, consisting of spatially ordered progenitor domains separated by sharp boundaries. At the other end of the D-V axis, Bone Morphogenetic Proteins (BMPs) produced by the roof plate regulate patterning of the dorsal progenitor domains ^23–28^.

However, both morphogens play a limited role in specifying the identity of neural progenitors at more intermediate positions of the D-V axis^29^. These constitute approximately half the cell types in this system and, like dorsal and ventral progenitors, are organized into discrete stripe-like domains^21,25,30–32^. Genetic and pharmacological perturbation of Shh signaling has shown that, while essential for specifying the three ventral progenitors closest to the source, it is dispensable for specifying more intermediate progenitors such as p0 and p1 cells ^33–35^ (Figure 1A). Similarly, BMP signaling appears to only be strictly required for the specification or differentiation of the two or three most dorsal progenitor cell types ^23,25,28^. Thus, alternative mechanisms must exist to pattern the intermediate regions of the neural tube, but these have remained elusive.

**Figure 1.**
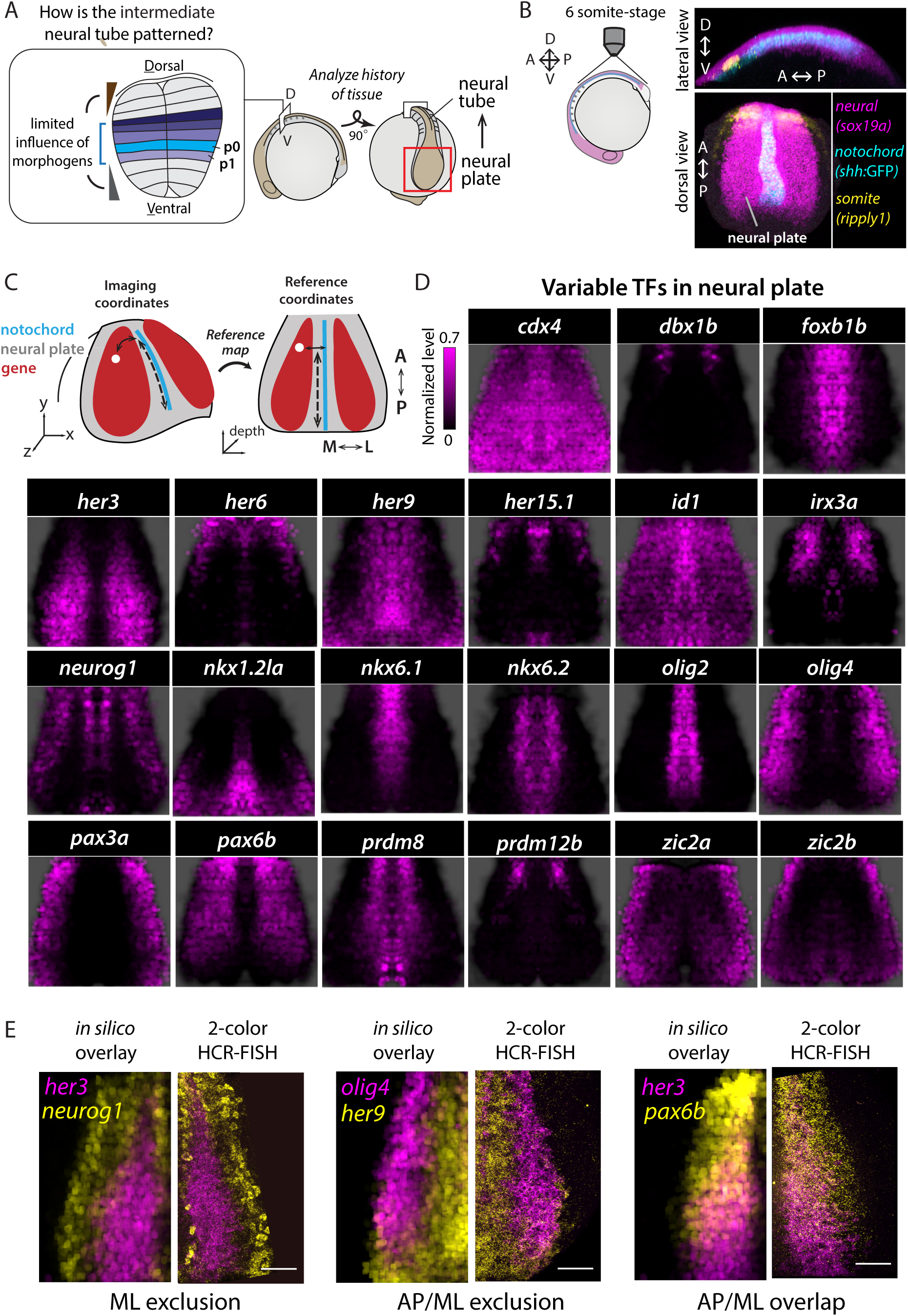
Reference-mapping reveals extensive neural plate patterning. (A) (Inset, schematic) Intermediate levels of the neural tube are patterned into multiple neural progenitor domains (colored stripes in transverse section, middle). Dorsal and ventral morphogen gradients (brown, gray triangles) do not direct patterning in this region. (Right, schematic) The neural tube is generated continuously from the neural plate as the embryonic axis extends. Therefore, at any given time, the neural plate contains cells at earlier developmental stages, representing a historical state of the tissue. (B) (Left) Schematic showing approximate zebrafish embryo orientation used to visualize the neural plate at the 6 somite stage (ss). (Right) 3D rendering of posterior 6 ss embryo expressing *shh:GFP* (cyan), with HCR RNA-FISH labeling of *sox19a* (magenta) and *ripply1* (yellow) expression, and imaged as in (A). “A” = Anterior; “P” = Posterior; “D” = Dorsal, “V” = Ventral. (C) Schematic showing correspondence between imaging coordinates and reference-based coordinates during reference-based mapping. “M” = Medial; “L” = Lateral. See also Supplementary Figure 1B, C. (D) Reference-maps showing mRNA expression (magenta) of the indicated transcription factors, measured using HCR RNA-FISH. The black background corresponds to the neural plate, co-labeled in each measurement using *sox19a*. Each map depicts the average of reference-mapped HCR RNA-FISH signal across n = 3 embryos, each symmetrized and normalized to the maximal detected signal within the image. See also Supplementary Figure 1E-G. (E) Comparison between *in silico* overlays of the reference maps for the indicated genes (left half of each split panel) and representative examples of their co-expression within a single embryo (right half of each split panel, maximum intensity projection), measured using two-color HCR RNA-FISH. Scale bar indicates 50 μm.

We hypothesized that the developmental history of the neural tube could impact patterning. The neural tube develops from the neural plate, a tissue that experiences a distinct and complex signaling environment due to its proximity to multiple signaling centers such as the notochord, tailbud, neural plate border, and newly-formed somites. The signals produced by these tissues, such as Wnts and FGFs, are thought to primarily regulate neural competence, anterior-posterior regionalization, and timing of neural differentiation^36–40^. Intriguingly, some neural factors are known to show patterned expression at this stage, suggesting that these signals could also provide positional information^37,41–44^. Nevertheless, a systematic analysis of neural plate signaling, patterning and effect on downstream fates has been missing.

Here we generated an integrated atlas of signaling, transcription factor states and cellular motion in the zebrafish neural plate and used it to identify the basis for intermediate neural tube patterning. We found that the juxtaposition of tailbud-derived Wnt signaling and lateral neural plate factors generates the positional information to establish precursor states that specifically give rise to p0 or p1 neural progenitors. Convergent-extension movements move precursor cells relative to Wnt and FGF signaling domains, altering their signaling exposure to induce progenitor specification. We show that, in the case of p0 progenitors, these movements also shape the domain and regulate the balance between specification and precursor maintenance. This morphogenesis-mediated patterning mechanism thus operates alongside conventional morphogen-mediated patterning to generate cell type diversity and organize the developing neural tube.

## Results

### An atlas of transcription factor expression in the neural plate reveals extensive patterning

Previous work on dorsal-ventral (D-V) patterning of the neural tube suggests that neural progenitors in the intermediate region are specified even in the absence of signaling from the ventral morphogen, Shh^30,31,33–35^ (Figure 1A). We first confirmed this by analyzing the expression of markers of intermediate p0 and p1 domains in zebrafish embryos after treatment with an Shh pathway inhibitor, from the onset of gastrulation (50% epiboly) until the 18 somite-stage. Under these conditions, p0 and p1 markers continue to be expressed in a patterned manner in the posterior neural tube, but a marker of the ventral pMN domain is completely extinguished (Supplementary Figure 1A).

We asked whether patterning might begin earlier, in the preceding neural plate stage, which continuously generates the neural tube as the body axis extends (Figure 1A). We sought to capture patterning in the neural plate by systematically analyzing the spatial expression of heterogeneous transcription factors (TFs). To achieve this, we developed a reference-based mapping approach to integrate spatial measurements made in different stage-matched embryos (Figure 1B, C). For reference-mapping, measurements were performed in transgenic embryos expressing a *shha:GFP* reporter^45^, used to label the notochord, and a ubiquitous membrane-localized mNeonGreen protein, which labels cell boundaries. The expression of the pan-neural TF *sox19a* was used to identify the neural plate (Supplementary Figure 1B, C, Methods). Each cell was individually segmented based on its membrane fluorescence and the following new coordinates were calculated relative to the notochord reference: (1) medio-lateral (M-L) distance from the notochord midline, (2) antero-posterior (A-P) distance from the posterior end of the notochord, and (3) depth from the surface (Figure 1C, Methods). These coordinates are contained within the frame of reference of the embryo and therefore should be more robust to variations in the orientation of embryos during imaging. We verified this using a broadly expressed neural plate gene, *olig4*^44^, and found that its reference-based expression maps remained relatively unchanged despite changes in the orientation of imaging (Supplementary Figure 1E-G).

Next, to identify heterogeneous neural plate TFs in an unbiased manner we analyzed previous single-cell transcriptomic data from zebrafish embryos^46^, focusing on TFs in the posterior neural tissue that are highly variable, display non-redundant patterns across cells and are expressed above a threshold value (Methods). This analysis returned 28 TFs. We supplemented this list with four known marker genes for early and intermediate spinal cord fates (*nkx1.2la*, *dbx1b, prdm12b,* and *irx3a*). We then used dual-color HCR RNA-FISH^47^ to measure the expression of each of these genes along with *sox19a* as a neural marker. We performed these measurements in 6 somite-stage (ss) embryos; at this stage, most of the neural plate is a monolayer (Supplementary Figure 1C) and could be analyzed as a pseudo-2D tissue, simplifying subsequent analyses. Moreover, photo-labeling cells in embryos expressing the photoconvertible protein kikGR^48^ showed that the 6 ss neural plate contributes to most of the spinal cord (somite levels 9-27) (Supplementary Figure 1D, Methods).

Our analysis revealed a highly patterned neural plate (Figure 1D, Supplementary Figure 2A-C). Out of our list of 32 candidate TFs, we could detect expression of 21 TFs within the neural plate. Other TFs were either expressed primarily in non-neural regions (*tfap2a, nr2f5*) or in the tailbud (*cdx1a)*, or were detected at very low levels, consistent with their low expression in previous *in situ* analyses (ZFIN) as well as a recent spatial transcriptomic dataset^49^. Comparing TF expression after reference-mapping showed that most TFs display unique spatial patterns, but together their expression covered the entire neural plate, indicating that our TF selection procedure captures spatial heterogeneity across the neural plate. Notably, the unique patterns observed for most TFs suggests diverse upstream regulation and substantial positional information at the neural plate stage.

Interestingly, these maps showed that significant D-V patterning of the future neural tube occurs in the neural plate but with differences between ventral, intermediate and dorsal regions in the timing of patterning. Markers for ventral neural progenitor fates such as *olig2, nkx6.1,* and *nkx6.2* are patterned even at the most posterior levels of the neural plate (Figure 1C, Supplementary Figure 3A). Since the neural plate morphs into the neural tube progressively from anterior to posterior as the embryonic axis extends, these posterior levels will not join the neural tube until the end of segmentation and axis extension (∼24 hpf). Yet, ventral patterning appears to be established well in advance of this transition. On the other hand, markers for intermediate neural progenitor cell types such as *irx3a, prdm12b* and *dbx1b* appear at more anterior levels of the neural plate, presumably closer in time to the neural tube transition. Moreover, while some broad markers of dorsal progenitor fates such as *pax3a* and *olig4* are expressed in the neural plate, more specific markers such as *ascl1b* are only detected in the neural tube (Supplementary Figure 3C).

Together, these data showed that reference-based mapping could be used to aggregate spatial information across embryos and revealed that the neural plate is a richly patterned tissue even before neural tube formation. D-V patterning begins in the neural plate but proceeds asynchronously in ventral, intermediate and dorsal sections of the future D-V axis, suggesting different underlying regulatory mechanisms.

### The neural plate comprises multiple TF states organized by a combination of signaling inputs

The diverse spatial patterns of neural TFs indicated that they are co-expressed in several distinct combinations. We found that reference maps could be used to reliably infer TF co-expression: comparing the *in silico* overlays of maps with directly measured co-expression for selected pairs of genes showed that reference mapping could infer co-expression patterns along both the medio-lateral (ML) and antero-posterior (AP) axes (Figure 1E, Supplementary Figure 2D).

We used *in silico* co-expression analysis to ask how many distinct regulatory states, defined as specific combinations of TFs, could be identified in the neural plate. We first smoothed the reference maps by agglomerating pixels into approximately cell-sized ‘superpixels’. Each superpixel could be assigned an expression level value for every TF based on its intensity in the corresponding TF maps. We then analyzed the gene expression states represented by these superpixels using *k-*means clustering (Figure 2A, Methods). This showed that the neural plate could be resolved into at least 16 co-expression states (Methods, Figure 2B, Supplementary Figure 4A). Visualizing the spatial locations of cluster labels revealed a grid-like organization of TF states, indicating that detailed antero-posterior and medio-lateral spatial information is encoded throughout the plate (Figure 2C, D). Notably, even the posterior region of the plate, which does not contribute to the neural tube until the end of axis extension (Supplementary Figure 1D), is patterned into distinct TF states (#13-16) at this stage.

**Figure 2.**
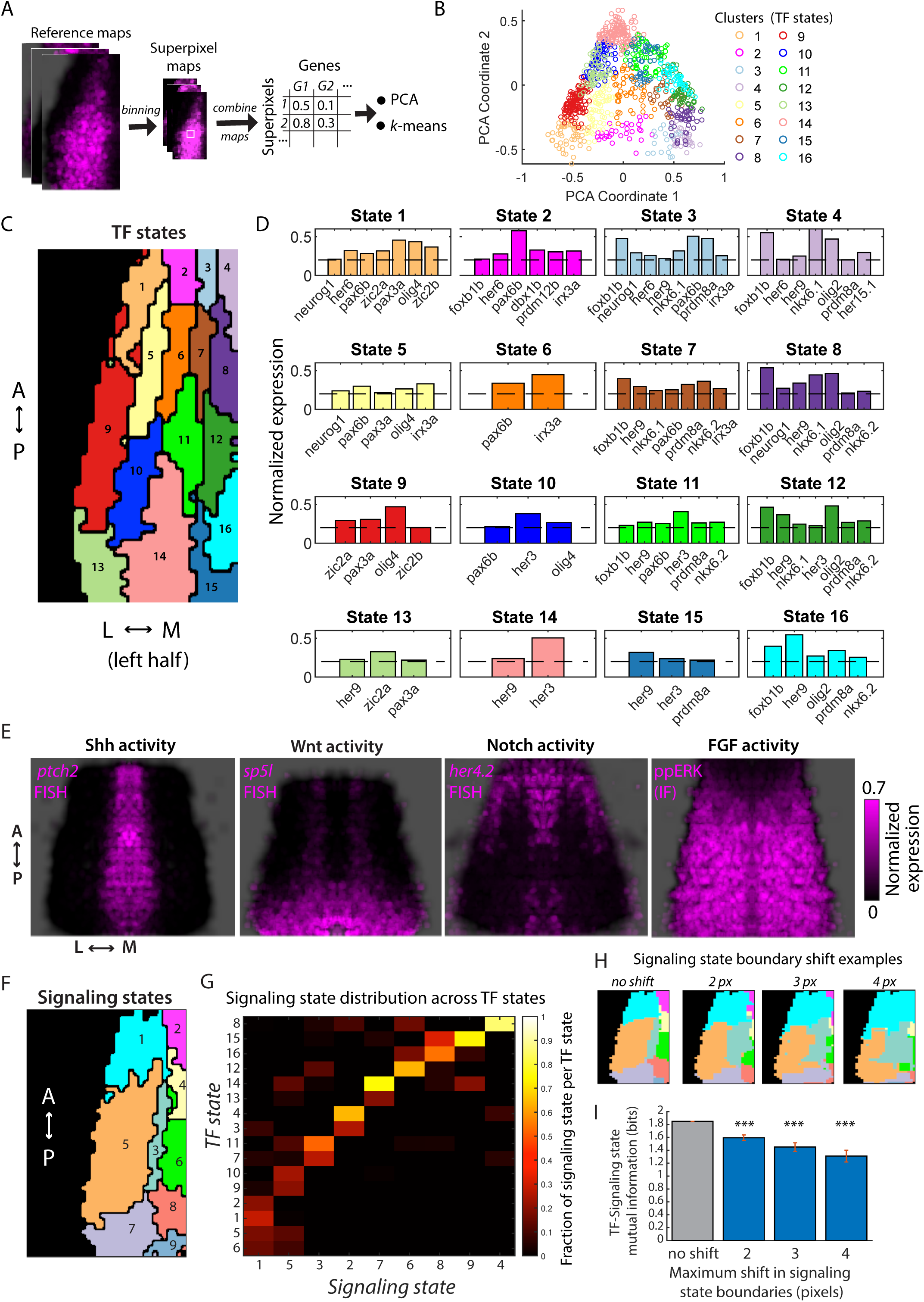
Neural plate patterning represents multiple transcription factor states that correspond to combinatorial signaling states. (A) Schematic of analysis workflow. Pixels in transcription factor (TF) reference-maps were binned to generate ‘superpixel’ maps, which have lower resolution but smoother TF expression. Expression in each superpixel was tabulated across TF maps to generate a superpixel vs. expression level matrix. Principal component analysis (PCA) and *k-*means clustering were applied to this data to analyze the spatial distribution of TFs. (B) Scatter plot showing principal component values for superpixels, colored by the clusters they are assigned to after *k*-means clustering. (C) Map of TF-state labels for superpixels after *k-*means clustering. (D) Mean expression levels of indicated TFs within the individual states shown in (C). Only TFs with mean expression above a threshold (dashed line) are shown. (E) Reference-maps of indicated signaling pathway targets. Each map depicts the average of reference-mapped signal across n = 3 embryos, symmetrized and normalized to the maximal detected signal within the image. (F) Map of signaling-state labels after *k-*means clustering of superpixels based on their levels of signaling pathway targets. See also Supplementary Figure 4D, E. (G) Heatmap showing how signaling states are distributed across TF states. Each column shows the fractional distribution of the corresponding signaling state across all the TF states. Note how most signaling states are specific to a subset of TF states. (H, I) Changes in mutual information between signaling and TF states when signaling state boundaries are perturbated *in silico*. (H) Representative examples of signaling states after boundaries are modified within the number of pixels indicated on top. (I) Mutual information between perturbed signaling states and TF states (blue bars) compared to that between the original (unperturbed) signaling states and TF states (grey bar). *** indicates *P* < 0.01 for the comparison with unperturbed mutual information, by Dunnett’s test.

Next, to understand the basis of patterning, we analyzed signaling in the neural plate. Previous work has suggested that ligands for five pathways – FGF, Wnt, Notch, Shh, and BMP - are expressed in the neural plate and tailbud region^36,50–52^. We systematically analyzed the activity of these pathways using either HCR RNA-FISH for well-established target genes (Wnt, Notch, Shh) or immunofluorescence to detect phosphorylated pathway effectors (FGF, BMP). As with transcription factor analyses, these measurements were made along with HCR RNA-FISH for the neural marker *sox19a* in *shha:GFP* transgenic embryos, enabling reference mapping.

We found that there was no discernible BMP activity (phosphoryated-Smad1/5/8 immunofluorescence) in the neural plate (Supplementary Figure 4B), but other pathways show patterned activity (Figure 2E). In particular, in addition to Shh signaling along the midline, we observed the Wnt target *sp5l* ^51^ in the posterior, the Notch target *her4.2*^53^ in the anterior, and a posterior to anterior gradient of the FGF target ppERK. Strikingly, the activity region of each signaling pathway corresponded to particular TF states, with shared boundaries, suggesting that patterns of TF expression could be defined by these signals (Supplementary Figure 4C).

Importantly, following a similar procedure as above for TF state identification, the neural plate could be subdivided into at least nine distinct signaling states defined by specific combinations of the four signals (Figure 2F, Supplementary Figure 4D, E). To understand the extent to which these signaling states could explain the observed pattern of TF states, we calculated the mutual information (MI) between signaling and TF states based on their fractional overlap (Methods). We observed that most signaling states corresponded primarily to one or two TF states, resulting in an MI value of 1.78 bits (Figure 2G). This means that knowing the signaling state of a cell narrows its plausible TF states by 2^1.78^ = ∼3.4-fold on average, indicating that the combination of the four signals could explain a substantial fraction of TF state diversity. In contrast, knowing the activity of a single signal such as Shh only narrows plausible TF states by ∼1.3-fold, a small (30%) increase over a random guess about the TF state (Supplementary Figure 4F, G). We note that this analysis does not consider possible differences in the effects of different levels of signaling, which would effectively increase the number of signaling states, but highlights the increased explanatory power of multiple signals in this context.

To further assess how well the observed configuration of signaling states explains TF state patterning, we analyzed how MI changes when signaling states are perturbed *in silico*. We took two approaches to computationally modify signaling states: First, we changed the boundaries of signaling states to varying extents by randomly reassigning pixels near the boundaries of each state to neighboring states and found that MI decreases consistently upon boundary changes (Figure 2H, I, Methods). Second, we allowed states to expand by reassigning to them broader pixel regions of their neighbors and again found that MI decreases with increasing deviation in the state map from the original (Supplementary Figure 4H, I). Both analyses suggest that the observed shapes and positions of the signaling states carries the most information about the observed pattern of TF states.

Together, these results showed that the neural plate is partitioned into multiple regulatory states marked by distinct combinations of TFs and suggested that four signaling pathways - Wnt, FGF, Shh and Notch – act in combination to pattern these TF states. We note that the averaging of multiple embryos to generate the reference map and the subsequent downsampling for clustering analysis reduces local heterogeneity in gene expression, such as in the case of salt and pepper expression of *neurog1* (Supplementary Figure 4J). These analyses therefore likely underestimate the number of TF and signaling states. Nevertheless, they capture broad differences across subregions of the neural plate.

### Intermediate neural progenitors arise from pre-patterned TF states in the posterior neural plate

As noted above, p0 and p1 neural progenitors, which eventually occupy intermediate D-V positions in the neural tube, in fact appear in the anterior of the neural plate, based on expression of their respective marker TFs *dbx1b* and *prdm12b* (Figure 1D). Thus, they appear to be specified in the context of the signals and states present in the neural plate. We next focused on identifying where they originate and how they are specified.

As the embryonic axis extends, the neural plate undergoes convergent-extension. Consequently, cells in the anterior neural plate join the neural tube, while other cells remain in the plate but occupy relatively more anterior positions (Supplementary Figure 5A). This suggested that cells might progressively transition from posterior TF states to more anterior states. We hypothesized that patterns of cell movement in the neural plate could be used to infer transitions between TF states. We therefore monitored the neural plate between 2 somite-stage (ss) to 6 somite-stage using timelapse imaging of a nuclear-localized H2B-mCherry fluorescent protein controlled by the *sox19a* promoter (Figure 3A, Supplementary Movie 1, Methods). These embryos also expressed the notochord marker (*shha:GFP*). Following automated tracking analysis, cell nuclei could be reference-mapped at each time point to generate a map of cell motion relative to the posterior end of the notochord (Figure 3B-C, Methods). This map revealed the average direction of cell motion at each point in the neural plate and confirmed that, as the axis elongates, cells move to relatively more anterior positions (further from the posterior end of the *shha:GFP* domain) while converging towards the midline.

**Figure 3.**
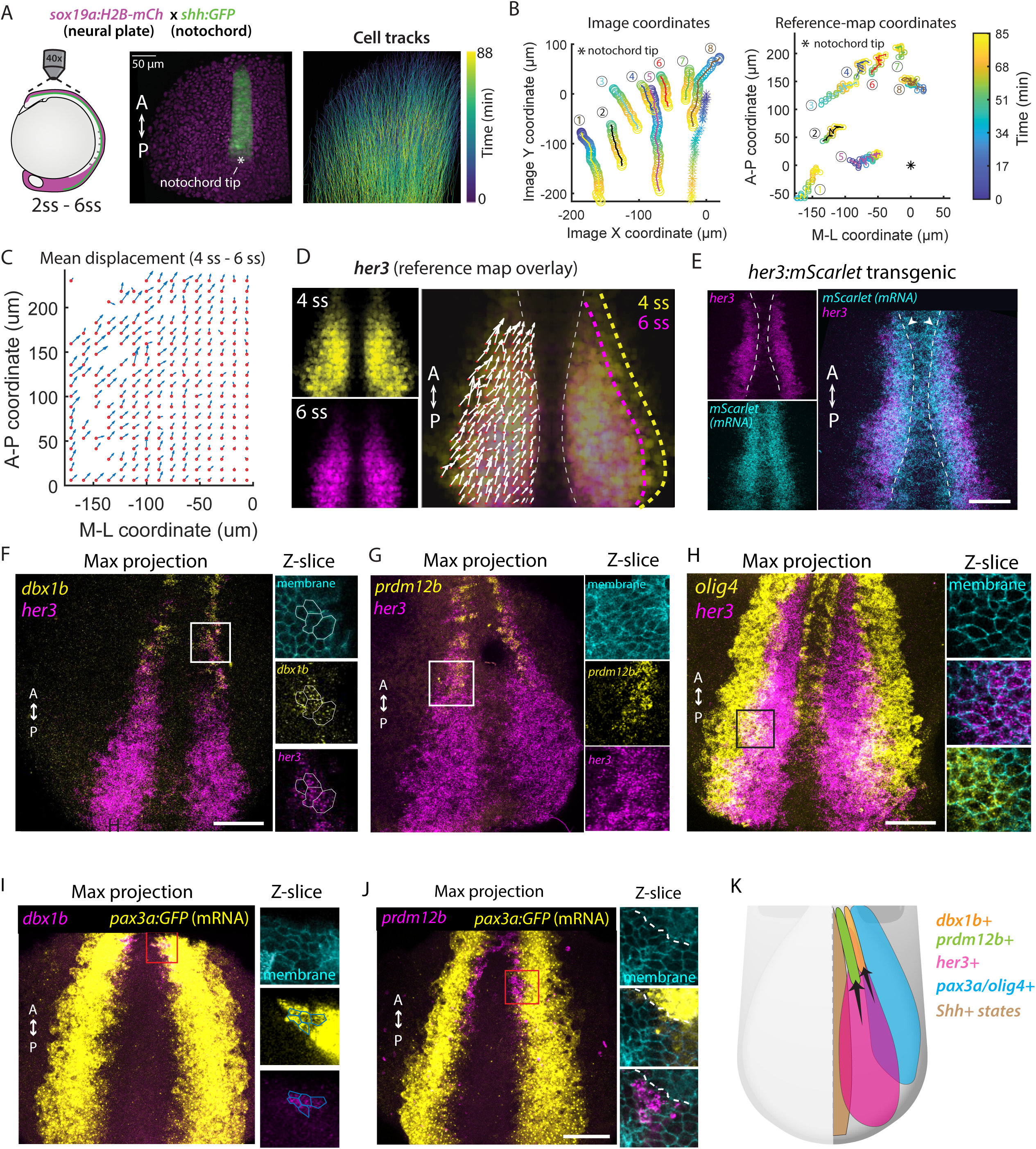
p0 and p1 neural progenitors arise from specific precursor states in the posterior neural plate. (A) (*Left*) Schematic showing region of *sox19a*:*H2B-mCherry*; *shh*:*GFP* double-transgenic embryo that was imaged to analyze cell movement in the neural plate. (*Middle*) 3D-rendered visualization of imaged area at 4 ss. Asterisk indicates the approximate position of the posterior tip of the notochord, annotated manually. “A” = Anterior; “P” = Posterior. (*Right*) Cell tracks indicating the position of cells at different timepoints during timelapse imaging. (B) Position over time for eight selected cells (circles linked by colored lines) and the notochord tip reference point (asterisk), in image coordinates (*left*) and after reference-mapping relative to the reference point (*right*). (C) Average displacement vectors (blue arrows) for cells in the neighborhood of the indicated positions (orange markers) between 4 somite stage (ss) and 6 ss. Positions and displacement vectors have been reference-mapped, and vectors from corresponding positions on the left and right sides of the embryo have been averaged. (D) Overlay of displacement vectors from (C) with reference maps of *her3* at 4 ss (yellow) and 6 ss (magenta). Yellow and magenta colored dashed lines indicate approximate lateral boundaries of *her3* expression at 4 ss and 6 ss, respectively. White dashed lines indicate approximate medial boundaries of *her3*. Note that displacement vectors outside *her3* expression regions are not shown. (E) Overlay of *her3* expression (magenta) and *her3*:*mScarlet-NLS* reporter expression (cyan), measured by HCR RNA-FISH. Images represent maximum intensity projections of 3D images. Dashed white line indicates approximate medial boundary of *her3* expression. Arrowheads indicate regions outside the *her3* boundary where *Scarlet-NLS* expression can be observed. Scale bar represents 50 *μ*m. (F) (*Left*) Representative example of *her3* (magenta) and *dbx1b* (yellow) expression, measured using HCR RNA-FISH, in a single transgenic embryo expressing a membrane-localized mNeongreen fluorescent protein (not shown). Maximum projection of 3D image is shown. White box indicates a region of *her3*, *dbx1b* co-expression, magnified on the right. (*Right*) A single Z-slice of the region outlined by the white box on left. Spectral channels for membrane mNeongreen fluorescence (cyan), *her3* expression (magenta) and *dbx1b* expression (yellow) are shown separately as well as together (‘merge’). Representative cells co-expressing *her3* (magenta) and *dbx1b* (yellow) expression are outlined. Scale bar represents 50 *μ*m. (G) (*Left*) Representative example of *her3* (magenta) and *prdm12b* (yellow) expression, measured using HCR RNA-FISH, in a single transgenic embryo expressing a membrane-localized mNeongreen fluorescent protein (not shown). Maximum projection of 3D image is shown. White box indicates a region of *her3*, *prdm12b* co-expression, magnified on the right. (*Right*) A single Z-slice of the region outlined by the white box on left. Spectral channels for membrane mNeongreen fluorescence (cyan), *her3* expression (magenta) and *prdm12b* expression (yellow) are shown. Scale bar represents 50 *μ*m. (H) (*Left*) Representative example of *olig4* (magenta) and *her3* (yellow) expression, measured using HCR RNA-FISH, in a single transgenic embryo expressing a membrane-localized mNeongreen fluorescent protein (not shown). Maximum projection of 3D image is shown. Black box shows a region of *olig4*, *her3* co-expression, magnified on the right. Scale bar represents 50 *μ*m. (*Right*) A single Z-slice of the region outlined by the black box on the left, showing membrane mNeongreen fluorescence (cyan), either separately (top), or overlaid with *her3* expression (*magenta*, middle) or *olig4* expression (*yellow*, bottom). (I) (*Left*) Representative example of *dbx1b* (magenta) and *pax3a:GFP* reporter expression (*yellow*), measured using HCR RNA-FISH, in a single transgenic embryo also expressing a membrane-localized mNeongreen fluorescent protein (not shown). Maximum projection of 3D image is shown. Black box indicates a region of *dbx1b, pax3a:GFP* co-expression, magnified on the right. Scale bar represents 50 *μ*m. (*Right*) A single Z-slice of the region outlined by the black box on left. Spectral channels for membrane mNeongreen fluorescence (cyan), *dbx1b* expression (magenta) and *pax3a:GFP* expression (yellow) are shown. Cells co-expressing *dbx1b* (magenta) and *GFP* (yellow) expression are outlined based on membrane fluorescence. (J) (*Left*) Representative example of *prdm12b* (magenta) and *pax3a:GFP* reporter expression (*yellow*), measured using HCR RNA-FISH, in a single transgenic embryo also expressing a membrane-localized mNeongreen fluorescent protein (not shown). Maximum projection of 3D image is shown. Black box indicates a region magnified on the right. Scale bar represents 50 *μ*m. (*Right*) A single Z-slice of the region outlined by the black box on left. Spectral channels for membrane mNeongreen fluorescence (cyan), *prdm12b* expression (magenta) and *pax3a:GFP* expression (yellow) are shown. Dashed line indicates medial boundary of *pax3a:GFP* expression. Note lack of *prdm12b* co-expression with *GFP.* Scale bar represents 50 *μ*m. (K) (*Schematic*) Model: *dbx1b*-expressing p0 cells (orange) emerge from cells co-expressing *her3* (magenta) and *olig4* (cyan), as well as *pax3a* (not shown). *prdm12b-*expression p1 cells (green) emerge from *her3* cells that lack *olig4* expression and Shh-activity.

Overlaying the map of cell motion with *dbx1b* and *prdm12b* maps suggested that these cell types originate from a lateral-posterior location (Supplementary Figure 5B). Inspecting TF expression patterns showed that *her3* occupies much of this region, implicating states expressing *her3* as likely sources (Supplementary Figure 5C). However, *her3* expression fades in the anterior, particularly at its boundary with *dbx1b* and *prdm12b* expression, suggesting a transition in cell state. Indeed, comparing *her3* expression at 4 ss and 6 ss to cell motion during this period showed that while the *her3* lateral boundary shifts in the direction of cell motion, the medial boundary does not shift much despite the flow of cells towards it (Figure 3D). This supported the idea that cells transition from *her3*+ to *her3*- at the anterior-medial boundary. Analyzing a long-lived reporter regulated by the *her3* promoter^43,54^ (*her3:mScarletNLS)* confirmed that *her3-* cells in the medial region previously expressed the gene (Figure 3E, Methods). More specifically, we performed dual-color HCR-FISH and could identify cells at the boundary that co-express *her3* with *dbx1b* or with *prdm12b*, likely corresponding to cells in transition (Figure 3F, G).

*her3* is expressed across multiple TF states (Figure 2D). We focused next on identifying the specific *her3+* state(s) that give rise to *dbx1b-* or *prdm12b-*expressing cells. Based on the long-lived her3 reporter, we observed that *dbx1b+* cells arise preferentially from the lateral boundary of the *her3* region (Supplementary Figure 5D). We therefore asked whether there were any distinctive features of this region that might bias cells towards the *dbx1b+* fate. Indeed, examining neural plate TF states indicated that cells in this region co-express *olig4* with *her3* (state #10 in Figure 2C, D), which could be verified using dual color HCR-FISH (Figure 3H). We also observed that *pax3a* and *olig4* show highly correlated expression in this region (Supplementary Figure 5E), suggesting that *olig4* and *pax3a* are co-expressed with *her3* at its posterior-lateral boundary.

These observations raised the possibility that *dbx1b+* cells in the anterior neural plate originate specifically from a *her3/olig4/pax3a* co-expressing state in the posterior neural plate. To test this, we first analyzed *pax3a* or *olig4* expression in *dbx1b+* cells but found that these genes are not co-expressed with *dbx1b* (Supplementary Figure 5F, G). We therefore hypothesized that, like *her3*, *pax3a* and *olig4* are expressed in *dbx1b* precursor cells but downregulated before *dbx1b* is induced. Indeed, when we fate-mapped pax3a-expressing cells using a long-lived fluorescent reporter (tgBAC(*pax3a:GFP*))^55^ we found that the reporter gene is co-expressed with *dbx1b,* supporting the model that precursor cells express *pax3a* (Figure 3I). On the other hand, the pax3a reporter is excluded from *prdm12b+* cells (Figure 3J). Thus, pax3a/olig4 expression defines the boundary between the precursors of p0 (*dbx1b*+) and p1 (*prdm12b*+) cells.

Finally, we note that the medial boundary of p1 cells coincides with the boundary of *ptch2* expression, which marks Shh activity (Supplementary Figure 5H). This suggests that the precursors of p1 cells likely correspond to the *her3*+ cells located between Shh-active states and olig4/pax3a+ states (state #14 in Figure 2C, D). Together, these data show that p0 cells (*dbx1b+*) and p1 cells (*prdm12b+*) are already pre-figured by specific TF states in the posterior neural plate. p0 cells derive from a *her3+* state co-expressing *pax3a* and *olig4,* while p1 cells derive from an adjacent *her3+* state that excludes these genes (Supplementary Figure 5I, Figure 4E). Notably, the precursor state markers *pax3a* and *olig4* are in fact key TFs that define dorsal neural tube progenitors (Supplementary Figure 3B), but appear to play a transient role in defining the p0 and p1 lineages.

**Figure 4.**
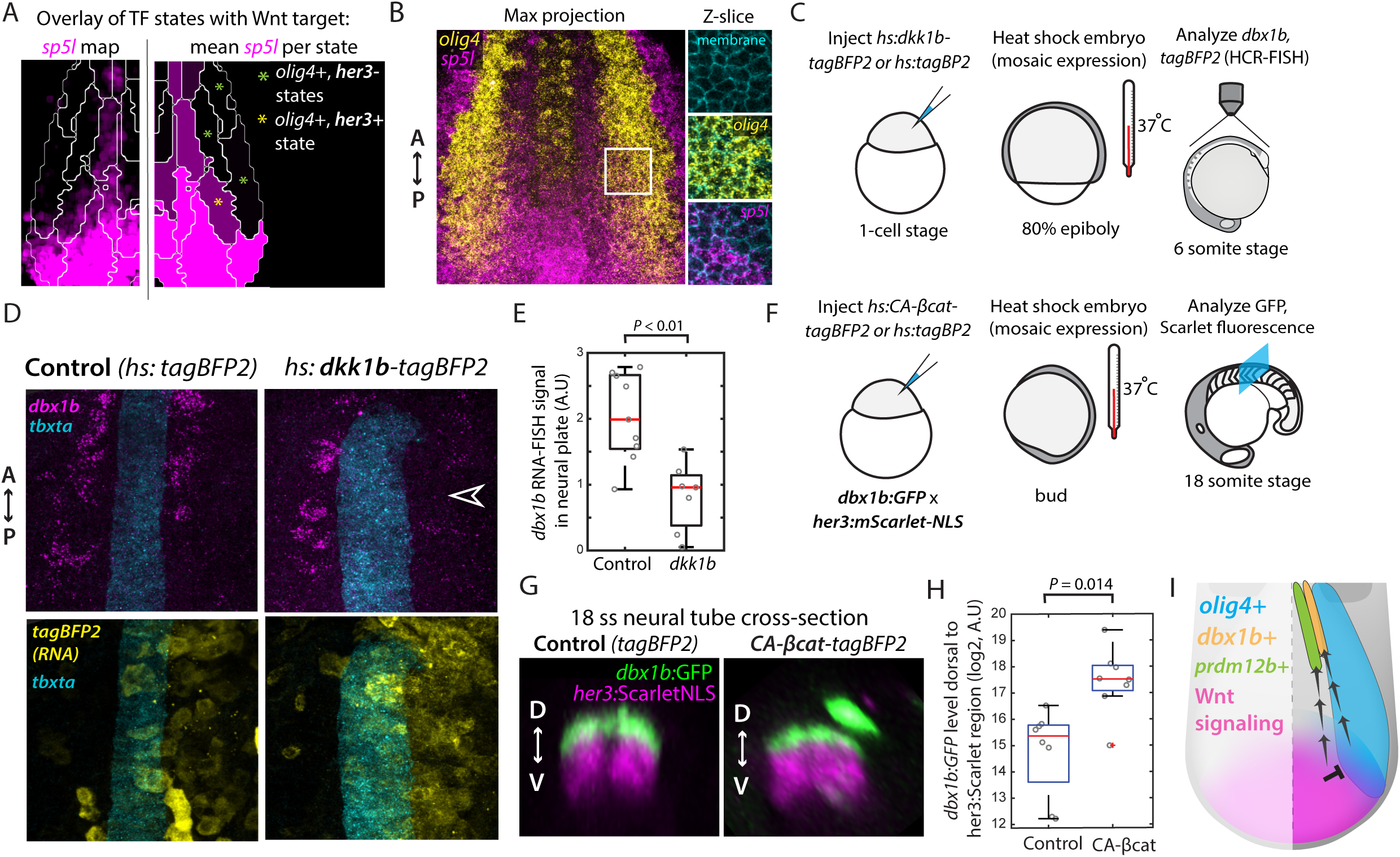
Interplay between Wnt signaling and *olig4* defines p0 and p1 precursors. (A) (*Left*) Outlines of TF states from Figure 2D overlaid on the reference map of *sp5l* expression (magenta, same as A but increased contrast). (*Right*) Mean level of *sp5l* expression within each TF state, calculated from the reference map of its expression. Green asterisks mark states expressing *olig4* but no *her3* (cf. Figure 2D), while the yellow asterisk marks the *dbx1b* precursor state, co-expressing *her3* with *olig4*. (B) (*Left*) Representative example of *sp5l* (magenta) and *olig4* (yellow) expression, measured using HCR RNA-FISH, within an individual embryo that also expresses a membrane-localized Neongreen fluorescent protein (not shown). Maximum projection of 3D image is shown. White box indicates a region of *olig4, sp5l* co-expression, magnified on the right. (*Right*) A single Z-slice of the region outlined by the white box on the left, showing membrane fluorescence (cyan) (top panel), overlaid with *sp5l* (middle panel) or *olig4* (bottom panel). (C-E) Effect of ectopic *dkk1b* (Wnt signaling inhibitor) expression on *dbx1b.* (D) Schematic of ectopic expression experiment. Embryos were injected at 1-cell stage with plasmids containing heat-shock inducible (‘hs’) dkk1b-tagBFP2 or tagBFP2. Genes were induced at the 80% epiboly stage, and expression of *dbx1b* was analyzed in the neural plate at 6 ss. (E) Representative images showing expression of *dbx1b* (top panels, magenta) in the anterior neural plate in embryos ectopically expressing *dkk1b-tagBFP2* (bottom right, yellow) or *tagBFP2* control (bottom left, yellow). *tbxta* expression (cyan) was used to mark the notochord. Expression was measured using HCR RNA-FISH. Empty arrowhead highlights reduction in *dbx1b* expression on the side of the embryo expressing higher levels of *dkk1b-tagBFP2*. (F) Boxplots showing total *dbx1b* expression, measured using HCR RNA-FISH as in (E), in embryos either ectopically expressing *dkk1b-tagBFP2* (n=7 embryos) or *tagBFP2* control (n=8 embryos). *P-*value calculated by Student’s t-test. (F-H) Effect of ectopic *CA-βcatenin* (‘CA-*βcat*’, Wnt signaling activator) expression on *dbx1b.* (G) Schematic of ectopic expression experiment. *dbx1b-GFP*; *her3:mScarlet-NLS* transgenic embryos were injected at 1-cell stage with plasmids containing heat-shock inducible (‘hs’) *CA-βcat-tagBFP2* or *tagBFP2*. Genes were induced at the bud stage, and *dbx1b:GFP* reporter fluorescence was analyzed in the neural tube at 18 ss. (H) Representative images showing *dbx1b-GFP* fluorescence in the neural tube of embryos expressing ectopic CA-*βcat*-tagBFP2 (right) or *tagBFP2* control (left). *her3:mScarlet-NLS* fluorescence (magenta) normally marks the lateral/dorsal limit of dbx1b:GFP and was used as a reference to identify ectopic GFP+ cells. (I) Boxplots showing total GFP fluorescence dorsal to the mScarlet-NLS+ region, in embryos either ectopically expressing *CA-βcat-tagBFP2* (n=8 embryos) or *tagBFP2* control (n=8 embryos). *P-* value calculated by Student’s t-test. See also Figure S6F. (I) Schematic showing that precursors of *dbx1b*+ cells (p0 cells) are defined by a region of low Wnt signaling (magenta) within the expression domain of *olig4* (cyan) in the posterior neural plate. On the other hand, precursors of *prdm12b*+ cells (p1 cells) are defined by a region of high Wnt signaling (magenta), which excludes *olig4* (cyan). Precursors become specified into corresponding *dbx1b*+ or *prdm12b*+ progenitors as cells move to a more anterior position in the neural plate.

### Precursor states are defined by tailbud-derived Wnt signaling

We next focused on the question of how the precursor states for p0 and p1 cells are defined. We observed that signaling states in the corresponding region of the neural plate are characterized by Wnt and FGF signaling. Indeed, Wnt activity (based on expression of the target gene *sp5l*) forms a gradient in the posterior-lateral region of the neural plate where p0/p1 precursor states are located (Figure 2E). Strikingly, comparing *sp5l* expression across TF states in this region indicated that the p0 precursor state (*olig4+/her3+*) state is distinguished from other *olig4+* states by its *sp5l* expression, albeit at low levels (Figure 4A). This was supported by HCR RNA-FISH for *olig4* and *sp5l*, which are generally expressed in complementary patterns (Supplementary Figure 6A), but co-expressed in a narrow region at the medial boundary of *olig4* (Figure 4B). We confirmed that this region corresponded to that of *her3/olig4* co-expression (Supplementary Figure 6B). Together, these data suggested that Wnt signaling within the *olig4* domain specifies the p0 precursor state.

To test the role of Wnt signaling, we reduced its activity by expressing the signaling inhibitor *dkk1b* under a heat-shock inducible promoter (Figure 4C, Methods). This led to a reduction in the co-expression of *her3* with *olig4* (Supplementary Figure 6C, D), as well as in the expression of the p0 marker *dbx1b* (Figure 4D, E). Thus, decreasing Wnt signaling shrinks the p0 precursor state and the specification of p0 cells downstream. We next asked whether ectopic Wnt signaling is sufficient for *dbx1b* specification. To test this, we expressed a constitutively active form of the Wnt pathway effector β-catenin (‘CA-βCat-tagBFP2’) in transgenic embryos that expressed a dbx1b reporter (TgBAC(*dbx1b:GFP)*^56^) (Figure 4F, Methods). These embryos also expressed the *her3:mScarletNLS* reporter, which normally encompasses the *dbx1b* expression domain (Supplementary Figure 5D). We subsequently analyzed GFP expression in the neural tube, where the dbx1b+ domain (i.e, p0 neural progenitors) is well-defined and ectopic cells can be identified and quantified more easily. We found that expression of CA-βCat-tagBFP2, but not a tagBFP2 control, led to ectopic GFP+ cells dorsal to the her3:mScarletNLS domain (Figure 4G, H). Since the dorsal neural tube arises from lateral regions of the neural plate, which is characterized by *olig4/pax3a* expression, this suggested conversion of *pax3a*+/*olig4+* cells outside the *her3* domain to *dbx1b*+ cells. To verify this, we performed timelapse imaging of the neural plate in *dbx1b:GFP*; *her3:mScarletNLS* embryos after CA-βCat-tagBFP2 induction and indeed could identify GFP+ cells that arise lateral to the *her3:mScarletNLS*+ region (Supplementary Figure 6E). We note that CA-βCat-tagBFP2 did not increase GFP+ cells in the ventral neural tube, which arises from the medial neural plate, supporting the view that Wnt signaling within the *pax3a/olig4*+ lateral region defines p0 precursor cells (Figure 4I)

Next, to understand how the p1 precursor state is defined, we asked what determines the medial boundary of *olig4* expression, which corresponds to the boundary between the p0 and p1 precursor states (Figure 3K). The complementary expression patterns of *olig4* and the Wnt target *sp5l* (Supplementary Figure 6A) suggested that there might be an antagonistic relationship between the two. We therefore hypothesized that high Wnt activity observed in the medial region inhibits *olig4* to limit its expression domain. Indeed, decreasing Wnt activity led to a shift in the *olig4* medial boundary towards the midline (Supplementary Figure 6G, H). Importantly, expression of the p1 marker *prdm12b* was diminished, suggesting that this medial encroachment of *olig4* shrinks the corresponding precursor state to reduce the number of p1 cells (Supplementary Figure 6I).

Together, these data support that Wnt signaling in the posterior neural plate acts in two ways to define both p0 and p1 precursor states. High Wnt activity near the midline blocks expression of the lateral neural plate factor *olig4* to create the p1 precursor state, while lower activity within the *olig4* domain creates the adjacent p0 precursor state. We note that Wnt signaling in the posterior neural plate likely derives from the tailbud region, outside the neural plate, based on expression of the ligands *wnt3a* and *wnt8a* (Supplementary Figure 6J). Thus, positional information to allocate p0/p1 precursors is generated from the juxtaposition of *olig4+* lateral neural plate with the tailbud.

### A convergent extension-based mechanism underlies intermediate progenitor patterning

The hallmark of D-V patterning in the neural tube is the stripe-like organization of progenitor domains. We next asked how such domains are derived from the precursor states allocated in the posterior neural plate, focusing on two questions: what controls the specification of progenitors from precursor cells in the anterior neural plate and the shape and size of the resulting progenitor domain.

Comparing reference maps for signaling and expression of p0 and p1 markers (*dbx1b* and *prdm12b*, respectively) showed that FGF signaling is active in much of the neural plate but drops off sharply in the anterior, where *dbx1b+* and *prdm12b*+ cells arise (Figure 5A, Supplementary Figure 7A). This suggested that a reduction in FGF signaling might lead to their specification from their respective precursors. Indeed, inhibiting FGF signaling using the small molecule SU5402 led to *dbx1b*+ and *prdm12b*+ cells appearing at more posterior positions in the neural plate (Figure 5B, C, Supplementary Figure 7B-D, Methods).

**Figure 5:**
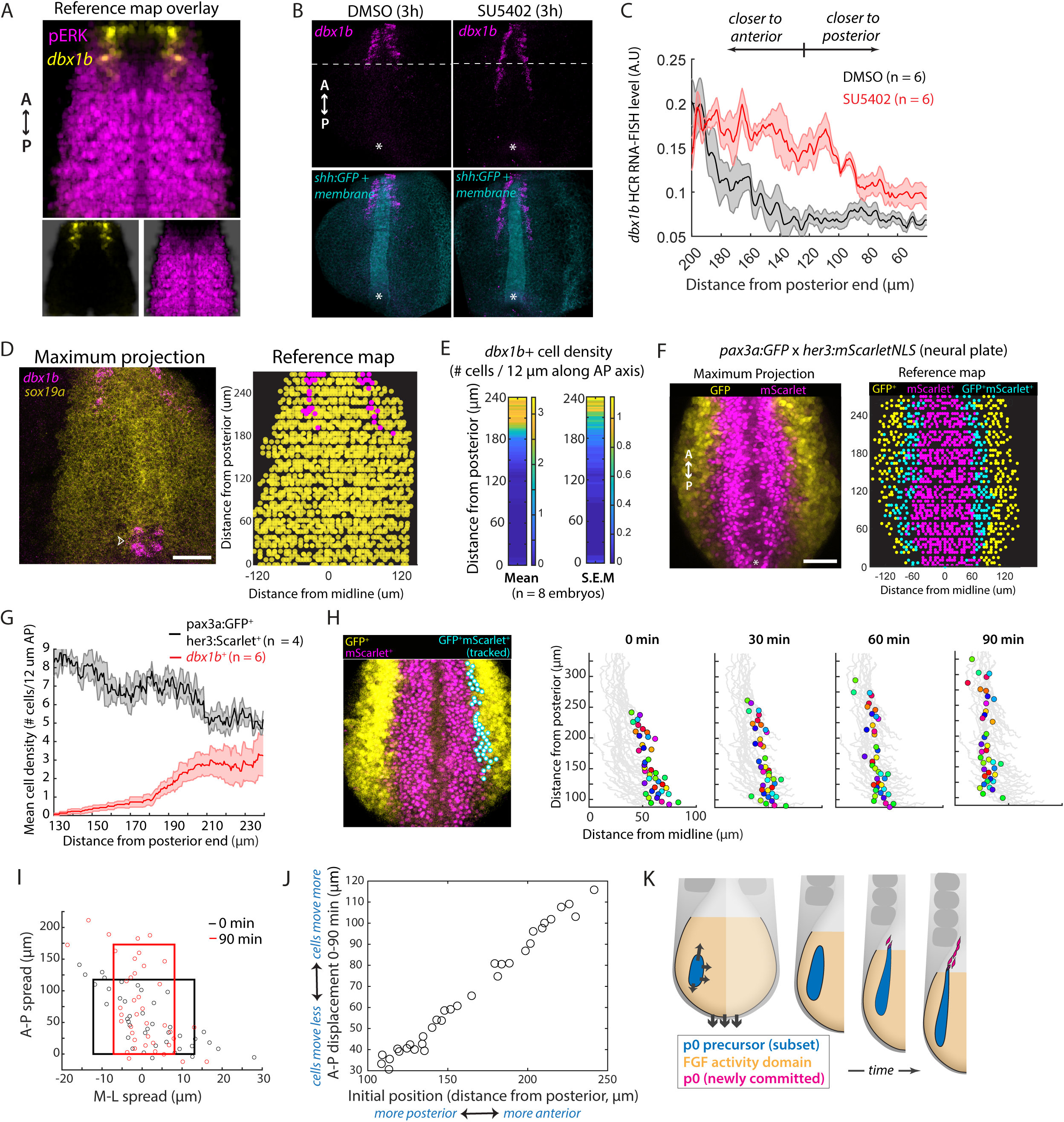
Convergent extension displaces precursors relative to FGF signaling to induce intermediate progenitor specification while shaping the domain. (A) Overlay of reference maps for *dbx1b* (yellow) and phosphorylated ERK (magenta) from Figure 1D and Figure 2E, respectively. (B) Representative images showing *dbx1b* expression (magenta), measured using HCR RNA-FISH, in transgenic embryos expressing *shh:GFP* and membrane-localized mNeongreen (cyan) after treatment with SU5402 or control (DMSO) for 3 hours. Shown are 3D rendered images from a dorsal perspective. The dashed line indicates the posterior extent of dbx1b in the DMSO-treated embryo, while the asterisk indicates the posterior end of the notochord based on *Shh:GFP* expression. (C) Axial profile of *dbx1b* expression in embryos treated with SU5402 (red) or DMSO control (black). Solid lines represent the median while shaded regions represent S.E.M. See also Supplementary Figure 7B, C. (D) (Left) Representative example showing expression of *dbx1b* and *sox19a*, measured using HCR RNA-FISH, in a transgenic embryo expressing the notochord marker *shh:GFP* and membrane-localized mNeongreen (not shown). Image is a maximum projection of a 3D image. Arrowhead indicates non-neural *dbx1b*+ cells ventral to the neural plate. (Right) Results of automatic identification and reference mapping of *dbx1b*+ cells (magenta) in the neural plate (marked by *sox19a* expression, yellow) based on the image shown on the left. Note that non-neural dbx1b cells are automatically excluded during reference-mapping analysis. Scale bar represents 50 *μ*m. (E) Heatmaps showing mean and S.E.M of the density of *dbx1b*+ cells at different axial levels in the neural plate. See also Supplementary Figure 7I. (F) (*Left*) Representative image showing GFP (yellow) and mScarlet-NLS (magenta) reporter fluorescence in the neural plate region in *pax3a:GFP; her3:mScarlet-NLS* double transgenic embryos that were injected with H2B-haloTag (not shown). Shown is a maximum projection of a 3D image. (*Right*) Results of automatic identification and reference-based mapping of cells expressing pax3a:GFP alone (yellow), her3:mScarlet-NLS alone (magenta), and both pax3a:GFP and her3:mScarlet-NLS (cyan) for the embryo shown on the left. Scale bar represents 50 *μ*m. (G) Axial profiles of cells co-expressing pax3a:GFP and her3:mScarlet-NLS (black) and *dbx1b* (red), based on reference maps such as those shown in (A) and (F). Solid lines represent the median while shaded regions represent S.E.M. See also Figure S7E. (H-I) Timelapse analysis of cells co-expressing pax3a:GFP and her3:mScarlet-NLS. (H) (*Left*) Maximum projection image of *pax3a:GFP*; *her3:mScarlet-NLS* double transgenic embryo used in timelapse imaging, highlighting cells co-expressing GFP and mScarlet-NLS (‘GFP^+^mScarlet^+^’) that were tracked over time (cyan, shown on the right). See also Figure S7F. (Right) Time course of reference-mapped GFP^+^mScarlet^+^ cell positions (colored circles). Gray lines show trajectories of movement over the timecourse period for these cells. (I) Comparison of aspect ratio of GFP^+^mScarlet^+^ cells at the beginning (black box) and end (red box) of the timecourse. Boxes span the 10^th^ – 90^th^ range in cell positions. (J) Schematic of proposed model for how p0 progenitors are continuously specified at a small but steady rate in the anterior neural plate. Any given subset of p0 precursors (blue region), initially located within the FGF activity domain (orange) in the neural plate, will extend anteriorly relative to the posterior end of the embryo due to convergent extension movements. The precursor domain constricts anteriorly due to disproportionate rates of movement. Anteriorly, cells will eventually begin leaving the FGF activity domain, inducing the p0 marker *dbx1b* (magenta). (K) (*Schematic*) Model for how p0 progenitors are continuously specified at a small but steady rate in the anterior neural plate. Any given subset of p0 precursors (blue region), initially located within the FGF activity domain (orange) in the neural plate, will extend anteriorly relative to the posterior end of the embryo due to convergent extension movements. The precursor domain constricts anteriorly due to disproportionate rates of movement. Anteriorly, cells will eventually begin leaving the FGF activity domain, inducing the p0 marker *dbx1b* (magenta).

What leads to a reduction in perceived FGF signaling within precursors? We observed that the relative A-P extent of the FGF signaling domain within the neural plate remains unchanged between two developmental stages (3 ss and 6 ss) (Supplementary Figure 7E, F). This is consistent with previous work showing that FGF signaling recedes with the posterior end of the embryo during axis elongation, because ligands are produced primarily in the tailbud region^57^. However, during this time, cells move to relatively more anterior positions within the neural plate (see Figure 3C) and thus will move out of the FGF signaling region. In the case of p0/p1 precursors, this cell movement relative to the FGF signaling field is expected to induce *dbx1b* or *prdm12b* expression respectively.

Finally, we asked what determines the size of intermediate progenitor domains in the neural tube, i.e., the number of cells specified at any given axial level. We focused on the p0 domain, since we could directly monitor both the precursor state (*her3+, pax3a+*) and the specified progenitors (*dbx1b*+) using the *her3:mScarlet-NLS*, *pax3a:GFP* and *dbx1b:GFP* reporters described above. First, we noted that the p0 domain constitutes a strikingly narrow stripe in the developing neural tube, comprising only 1-2 dbx1b+ cells on each side of the midline at any axial level (Supplementary Figure 7G, H). Consistent with this, quantifying the spatial density of nascent *dbx1b*+ cells in the anterior neural plate showed that there are only 3 ± 1 cells per unit cell length (along the A-P axis) on either side of the midline (Figure 5D, E, Supplementary Figure 7I). This presumably reflects a limited rate of p0 cell specification from precursors. What determines this specification rate? To address this, we first sought to quantify the number of precursor cells at different A-P levels in the neural plate, particularly at the FGF activity boundary where the cells become specified (*dbx1b*+). We used a *her3:mScarlet-NLS*, *pax3a:GFP* double reporter line to identify precursors (Figure 5F, Supplementary Figure 7J). Importantly, this line also enabled tracking of precursor cells in the anterior neural plate, after specification, because GFP and mScarlet reporter fluorescence persists in specified *dbx1b*+ cells despite loss of *pax3a* and *her3* expression (Figure 3F, I, Supplementary Figure 5D, G). Quantifying GFP^+^mScarlet^+^ cells showed that precursor density in the neural plate decreases from posterior to anterior, reaching a comparable density to *dbx1b*+ cells (5 precursor cells vs. 2-4 *dbx1b*+ cells per cell length) (Figure 5G). This suggested that the rate of p0 specification is limited by the density of precursors in the anterior neural plate.

Next, to better understand how the density of precursors is regulated along the A-P axis, we performed timelapse imaging in *her3:mScarlet-NLS*, *pax3a:GFP* embryos. An H2B-haloTag was used to label nuclei, enabling tracking of precursor cells (GFP^+^ Scarlet-NLS^+^) as well as reference-mapping (Figure 5H, Supplmentary Figure 7K, Supplementary Movie 2, Methods). We observed that the precursor pool constricts medially but extends anteriorly over time, consistent with convergent extension movements (Figure 5H, I). Importantly, the axial extension of the pool is uneven, with cells being displaced a distance proportional to their initial axial position (Figure 5J). Specifically, the most anterior precursor cells traversed over 100 *μ*m over the 90 min imaging period, while the most posterior cells traversed as little as 30 *μ*m. This has the effect of simultaneously reducing the spatial density of precursors located anteriorly, i.e. thinning the precursor domain to limit the rate of specification of *dbx1b*+ cells, while continuing to maintain precursors in the posterior.

Together, these data suggested a model in which convergent extension movements in the neural plate perform three functions: (1) they drive specification of progenitors, by moving corresponding precursor cells out of an FGF signaling domain, (2) they shape the p0 progenitor domain and control its size by constricting the pool of precursor cells as they shift to more anterior positions in the neural plate and (3) they help maintain precursors in the posterior through elongation of the body axis (Figure 5K). Together, these functions enable patterning of p0 progenitors into a narrow stripe-like domain across the neural tube.

## Discussion

Discovering mechanisms for tissue patterning in diverse contexts is a fundamental goal in developmental biology. Here we identified a tissue-morphogenesis driven mechanism that patterns the intermediate levels of the posterior neural tube, which gives rise to the spinal cord. We find that, preceding neural tube formation, the neural plate is pre-patterned into multiple transcription factor states, which correspond to distinct regions of signaling (Figure 1, 2). Cells transit through these states as they undergo morphogenetic movements that alter the signals they experience (Figure 3-5). In particular, p0 and p1 neural progenitors, which occupy the intermediate neural tube, arise from specific precursor states that are established in the posterior neural plate through tailbud-derived Wnt signaling (Figure 4). Convergent extension movements enable continuous p0 and p1 progenitor specification from these precursors by effectively extruding them at a limited rate out of an FGF signaling domain (Figure 5).

While multiple factors act together to enable patterning in this system, neural plate morphogenesis is a central player, performing two key functions. First, it drives transitions in signaling, by moving cells relative to Wnt and FGF signaling domains. In particular, this enables the transient co-option of tailbud Wnt signaling (primarily associated with regulating body elongation and neuromesodermal progenitors^58–60^) to create p0/p1 precursor states by antagonizing lateral neural plate TFs like *olig4* (Figure 5I). Critically, tailbud Wnt signaling is known to inhibit neural differentiation^60–62^; cell movement out of this region thus permits downstream fate specification. Second, convergent-extension movements regulate the relative size of the p0 progenitor domain by constricting the precursor population as cells move anteriorly out of the FGF signaling domain (Figure 5K). Recent work has shown that morphogenetic movements also regulate patterning in the presomitic mesoderm by controlling signal exposure^63,64^, suggesting that this could be more broadly utilized patterning strategy.

Remarkably, the end result of morphogenesis-mediated patterning of intermediate progenitors, described above, and Shh-mediated patterning of ventral progenitors^14^, is similar: both result in stripe-like organization of progenitor domains in the neural tube (Supplementary Figure 1A). However, these mechanisms differ in key respects. First, the source of positional information is distinct. In the case of ventral patterning, this is supplied by a single factor, the Shh morphogen gradient. In the case of intermediate patterning, on the other hand, the positional information to establish p0/p1 precursor states comes from the interplay between Wnt signaling and lateral neural plate pre-patterning (Figure 4). Second, determination of progenitor identity and organization of progenitors into stripe-like domains are more tightly coupled in ventral patterning than in intermediate patterning. Ventral patterning occurs in a single step, with a single gene-regulatory network interpreting the Shh gradient to concurrently specify progenitor identity, determine relative proportions and establish domain boundaries^14,22^, though adhesion-based cell sorting corrects patterning errors^18,65^. In contrast, intermediate patterning proceeds through multiple steps. Progenitor identity is determined first, through the establishment of precursor states. Subsequently, convergent-extension and FGF signaling act together to regulate the rate of specification to control progenitor proportions. Finally, cell sorting likely helps establish domain boundaries after specification since it has been demonstrated that p0 cells sort through relatively strong homotypic adhesion^65^.

Our model supports and extends key observations from previous work related to intermediate progenitor specification. It explains why Shh and BMP signaling are largely dispensable for p0/p1 specification^29,30,34,35,66^. On the other hand, it is consistent with reports that retinoic acid (RA) signaling is important for p0 specification^67^, since a primary role for RA in the neural tissue is the attenuation of FGF signaling. Moreover, the model helps clarify why p0 and p1 fates are still specified in organoid models that are exposed only to transient Wnt and FGF activation^29,66,68^. Our findings that p0 cells arise from the pax3a-expressing lateral neural plate is consistent with work tracking Pax3 expression and function in the mouse neural tube^29,69^. Interestingly, this prior work shows that Pax3 in fact has a repressive effect on p0 specification and therefore must be inactivated for p0 cells to emerge. Our results suggest that Wnt signaling, which is a hallmark of the p0 precursor state (Figure 4B), could counteract pax3a for this purpose.

Finally, we note that patterned expression of TFs in the posterior neural plate has been evident since early pioneering studies of neural induction and dorsal-ventral patterning ^26,39,40,43,70,71^. Yet, the extent of patterning, and its regulation and impact on downstream fate specification has been unclear. The systems-level approach we undertook in this work, integrating systematic analyses of TFs, signaling and tissue dynamics, was crucial to elucidating these aspects of spinal cord patterning. Similar analyses have been applied recently to elucidate the relationship between signaling dynamics and mesodermal fate specification in zebrafish^64,72^. We anticipate that such approaches, with the incorporation of newer techniques for unbiased multimodal characterization of cell state^73–75^ and tissue dynamics ^6,76^ *in vivo*, will provide new insights and open up fundamentally new perspectives in understanding the organization of the spinal cord and other developmental systems.

## Acknowledgements

The authors thank members of the Megason and Lahav labs for helpful feedback. S.N. was the Philip O’Bryan Montgomery, Jr., MD, Fellow of the Damon Runyon Cancer Research Foundation (DRG-2339-18) and is supported by National Institute of Health grant 1K99GM151398. Research in the Megason Lab is supported by National Institutes of Health grant R01GM107733.

## Author Contributions

S.N, A.C and S.G.M designed experiments. S.N, A.C, B.Z.J, H.L and C.C performed experiments. D.E.W and T.Y-C.T generated key reagents and transgenic lines. S.N and S.G.M wrote the manuscript with input from all authors.

## Supplemental information

Document S1. Supplementary Figures 1–7 and supplemental references

Movie 1. Timelapse microscopy of neural plate in *sox19a:H2B-mCherry2* embryos, related to Figure 3

(Left) Timelapse of 3D-rendered confocal images showing dynamics of neural plate nuclei (purple) and the notochord (green) starting at 2 ss. (Right) Nuclei that were tracked in movie, colored by duration of tracks (blue = short, yellow/green = medium, red = long).

Movie 2. Timelapse microscopy of p0 precursors in *pax3a:GFP; her3:mScarletNLS* transgenic embryos, related to Figure 6

Timelapse of 3D-rendered confocal images showing dynamics of H2B-haloTag expressing nuclei (cyan) in the posterior neural plate region of the embryo. Note that H2B-haloTag is expressed broadly in the embryo, not just the neural plate. (Inset) Timelapse of 3D-rendered confocal images showing pax3a:GFP and her3:mScarletNLS expression in the same embryo. However, these images are acquired more infrequently (7.5 min intervals) compared to the H2B-haloTag images (1.5 min) to reduce photo-bleaching of reporters.

## Methods

### Zebrafish strains

Zebrafish (*Danio rerio*) were raised, maintained, and used for experiments as per protocols approved by the Harvard Medical Area Institutional Animal Care and Use Committee (HMA IACUC). AB and TL strain fish were used for all experiments. Juveniles and adults were maintained on a 14 h light / 10 h dark cycle at ∼28°C. Embryos were typically obtained from 4 mo – 18 mo old fish, and raised at 25 to 33°C.

The following new transgenic lines were generated for this study and are described below: Tg(*sox19a:H2B-mCherry2*) and Tg(*her3:mScarlet-NLS*). The following transgenic strains were also used and have been described previously: TgBAC(*dbx1b:GFP*)^nn11^ ^77^, TgBAC(*pax3a:GFP*)^i130^ ^55^, Tg(*shha:GFP*)^45^, Tg(*actb2:MA-2xmNeongreen*)^hm40^ The TgBAC(*pax3a:GFP*)^i130^ line was a kind gift from Dr. Phil Ingham.

### Generation of transgenic lines

#### Tg(sox19a:H2B-mCherry2)

An amplicon spanning a ∼6.7kb genomic sequence upstream of the predicted sox19a translation start site was PCR-amplified from a chromosome 5 BAC clone (CH73-378C22, CHORI-BACPAC) using primers 5-GCATAATCTAGCGCGAGTCC-3 and 5-CATGGCTGCCAACAGAAGT-3, Phusion polymerase, and the following cycling parameters: 98°C for 1 min, 35 cycles of 98°C for 10 sec, 64°C for 30 sec, and 72°C for 3 min, followed by 72°C for 5 min and a 4°C hold. The resulting amplicon was ligated into a pMTB vector backbone fragment (AddGene #112225) amplified with the same cycling parameters and primers 5-ACTTCTGTTGGCAGCCATGTCTAAAGGTGAAGAACTGTTCA-3 and 5-GGACTCGCGCTAGATTATGCGTAATGACTAGGCCCTCGAGC-3 by isothermal assembly to replace the original actb2 promoter with that of sox19a. Subsequently, reporter downstream of the promoter was replaced with an mCherry2 gene fused to histone 2B (H2B).

#### Tg(her3:Scarlet-NLS)

A ∼4.7 kb fragment of the *her3* promoter^43^ was amplified from a pBSceI-her3:GalTA plasmid ^54^ and cloned upstream of an mScarlet gene fused to an SV40 NLS sequence (PKKKRKV).

Transgenic founders were identified following the co-microinjection of single-cell stage zebrafish embryos with Tol2 mRNA and the resulting plasmid DNA.

### Confocal Microscopy

All imaging was performed on a Zeiss LSM980 laser scanning confocal microscope using a Plan-Apochromat 20X 1.0 NA, C-Apochromat 40X/1.1 NA, or a LD LCI Plan-Apochromat 40x/1.2 NA objective. A ∼300 μm x 300 μm x 100 μm region was imaged at a resolution in a typical experiment. For timelapse imaging, embryos were maintained at 25-28 C in egg water and mounted using a previously described procedure^18^ that allowed long-term imaging of the posterior neural plate. For imaging of fixed embryos, particularly for samples used to generate transcription factor and signaling activity maps, they were mounted in PDMS casts (10% cross-linker, with 1% Triton-X100) and embedded in Prolong Gold Antifade Mountant (ThermoFisher) prior to imaging using the LD LCI Plan-Apochromat 40x/1.2 NA immersed in glycerol.

### Reference-based mapping of fluorescent measurements in the neural plate

For reference-based mapping, measurements were made in transgenic embryos in which cells are uniformly labeled using a membrane- or nuclear-localized fluorescent protein, and the notochord is labeled using a *shha:GFP* reporter ^45^. Following confocal imaging, membrane-labeled cells were individually segmented using PlantSeg^78^, while nuclear-labeled cells were identified using Trackmate 7 ^79^. Custom MATLAB software was used to manually annotate the approximate dorsal midline of the notochord and to automatically calculate the outer surface of the embryo based on intensity thresholds.

Next, the sagittal plane of the embryo was calculated by using the anterior, posterior, and dorsal extrema of the annotated notochord midline. A mediolateral (ML) coordinate was assigned to each segmented cell/nucleus based on its distance from the sagittal plane, calculated along contours defined by the shape of the embryo surface. An anteroposterior (AP) coordinate was similarly calculated, based on the distance from the posterior tip of the annotated midline. Finally, the depth of the cell (D) was calculated based on its distance from the embryo surface, along the normal vector to the surface. These three coordinates constitute the ‘reference-based’ coordinates for each cell.

To generate the reference-map for a given measurement, cell segments were translated to their reference-based [AP, ML, D] coordinates within a new 3D image, with all pixel values within each segment assigned its mean signal value.

### Selection of transcription factors for neural plate characterization from single-cell data

A published transcriptomic dataset covering different stages of zebrafish embryogenesis^46^ was analyzed to identify potential spatially patterned transcription factors in the neural plate. SCANPY^80^ was used for analysis. Cells from 10 hours post fertilization (hpf) embryos annotated as ‘posterior neural tissue’ were organized into 10 Louvain clusters, and the ‘highly-variable’ function within SCANPY was used to identify the 200-most variable genes. These genes were then cross-referenced with annotated zebrafish transcription factors (AnimalTFdb 4.0^81^) to identify the most variable transcription factors (TFs). Next, TFs expressed above a maximum value of 0.3 within at least one Louvain cluster were identified. Genes that display redundant patterns of expression across clusters were then identified: for each pair of genes, their cross-correlation of mean expression across clusters was calculated, and genes were hierarchically clustered based on this. For sets of genes with median cross-correlation above a threshold value, only the highest expressed member was retained. This procedure yielded 28 TFs for further analysis. The expression of all identified TFs was then analyzed by HCR RNA-FISH. Some genes showed very low or undetectable HCR RNA-FISH signal within the neural plate and were not used further: *fosab, hnf1ba, junbb, meis3, pax6a,* and *id2a*.

### Identification of TF co-expression states and similarity

Reference-based maps were generated as 3D images at approximately the resolution of the original confocal image (0.59 um x 0.59 um x 0.7 um) but transformed into 2D by maximum projection along the depth axis. These maps were subsequently downsampled for analysis by binning pixels 10x10 (corresponding approximately to the size of a cell in the neural plate, 5 um diameter). For analysis of TF variation and identification of co-expression states, the map for each gene was transformed into a vector by concatenating rows in the image matrix. Principal Component Analysis and k-means clustering were performed on these map vectors.

Two metrics were used to determine the number of *k*-means clusters (i.e. TF states): (1) the silhouette score and (2) the range of expression values for each TF across clusters. The first metric decreases as clusters become more cohesive, while the second metric increases as superpixels with disparate values of TF expression are separated. We observed that the silhouette score decreased monotonically up to the maximum number of clusters analyze (20), while the expression range metric plateaued beyond 10 clusters. The value of 16 clusters chosen for downstream analyses represented a balance between low silhouette score (more cohesive clusters) and high expression range (better isolation of superpixels with the highest or lowest expression of genes).

For calculation of gene expression similarity, each cluster was analyzed as a vector in an 18-dimensional space, with dimensions corresponding to the 18 different genes displayed in Figure 2C and its value along each gene dimension corresponding to the mean expression of that gene within the cluster. Similarity was assessed using a cosine similarity distance.

### Identification of TF and signaling states and mutual information

Reference-based coordinates for cells were calculated in 3D, thereby generating 3D maps at approximately the resolution of the original confocal image (0.59 um x 0.59 um x 0.7 um), but transformed into 2D by maximum projection along the depth axis. These maps were subsequently downsampled for analysis by binning pixels 10x10 (corresponding approximately to the size of a cell in the neural plate, 5 um diameter). For analysis of TF variation and identification of co-expression states, the map for each gene was transformed into a vector by concatenating rows in the image matrix. Principal Component Analysis and k-means clustering were performed on these map vectors.

Two metrics were used to determine the number of *k*-means clusters (i.e. TF states): (1) the silhouette score and (2) the range of expression values for each TF across clusters. The first metric decreases as clusters become more cohesive, while the second metric increases as superpixels with disparate values of TF expression are separated. We observed that the silhouette score decreased monotonically up to the maximum number of clusters analyze (20), while the expression range metric plateaued beyond 10 clusters. The value of 16 clusters chosen for downstream analyses represented a balance between low silhouette score (more cohesive clusters) and high expression range (better isolation of superpixels with the highest or lowest expression of genes).

Mutual information between signaling and TF states was calculated as follows: for each signaling state, its distribution across TF states *T*_*i*_ was calculated based on the fractional overlap with each TF state yielding the conditional probability *P*(*T*_*i*_|*S*_*j*_). The mutual information *I* was calculated from the joint probability *P*(*T*_*i*_, *S*_*j*_) = *P*(*T*_*i*_|*S*_*j*_) ⋅ *P*(*S*_*j*_) using the standard definition 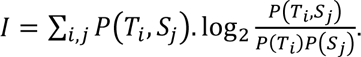

### in silico signaling state boundary perturbations

Signaling state boundaries were modified *in silico* using two approaches. First, pixels outside the boundary of each signaling state, within a. certain distance from the boundaries, were randomly assigned to the closest signaling state. Boundaries were then smoothed using standard morphological closing operations on the signaling state map. Second, for a given target signaling state, its area was increased by scaling factor, allowing it to encroach into neighboring states and thus modify the signaling state map. The mean fractional change in the areas of all signaling state was calculated to parametrize the degree of modification. Several modified signaling state maps were thus generated by systematically iterating through the target signaling state and scaling factors.

### Photolabeling of neural plate

Transgenic embryos expressing the photoconvertible fluorescent protein kikumeGR ^48^ were used to track the contributions of different regions of the neural plate to the spinal cord. Briefly, 6 ss embryos were mounted dorsally and rectangular regions in the neural plate were photoconverted using a 405 nm laser on a Zeiss 710 confocal microscope. Following labeling, embryos were incubated at 28 C until they reached 25 hpf. Embryos were then re-imaged from a lateral perspective to locate photo-converted cells within the posterior neural tube.

### in toto timelapse analysis of the neural plate

Microscopy: Transgenic embryos expressing *sox19a:H2B-mCherry2* and *shha:GFP* were dechorionated and mounted for dorsal imaging of the neural plate. Beginning at 2 ss (∼11 hpf), 3D confocal images were acquired at 1.5 min intervals using a 40x water-immersion objective in a temperature-controlled microscope setup maintained at 28 C.

Analysis: Following image acquisition, cell nuclei in the neural plate were tracked based on the H2B-mCherry2 label using the Trackmate v7 ^79^ plugin within ImageJ/Fiji ^82^. Briefly, nuclei were identified based on an intensity threshold and tracked using the Kalman tracker in Trackmate, followed by manual correction and curation of long tracks.

### Heat-shock induced Wnt perturbation

Heat-shock expression plasmids: *mtagBFP2, dkk1b-tagBFP2, or CA-β-catenin-tagBFP2* were cloned downstream of a *hsp70* promoter in vectors compatible with tol2-mediated transgenesis. *dkk1b-mtagBFP2* was generated by fusing mtagBFP2 at the C-terminus of *dkk1b* cDNA. *CA-β-catenin-tagBFP2* was generated by fusing mtagBFP2 at the C-terminus of *danio rerio ctnnb1* cDNA containing S>A mutations at the GSKβ 3 phosphorylation sites S33, S37, S41 and S45 (^83^.

Heat-shock induction: Embryos were co-injected at the 1-cell stage with one of the heat-shock expression plasmids and a plasmid containing the tol2 transposase. Embryos were raised to the heat-shock induction stage and subjected to 15 min exposure to 37 C to induce expression, before being returned to 25 or 28 C until time of analysis.

### Chemical perturbation of FGF signaling

SU5402 (SelleckChem, Catalog No.S7667) was stored at a stock concentration of 50 mM in DMSO. For perturbation experiments, bud-stage (10 hpf) embryos were dechorionated using Pronase (2 mg/ml) and then treated with 50 uM SU5402 in Danieau buffer. Treated embryos were raised at 28 C for 3h and subsequently washed 3 times with Danieau buffer prior to fixation.

### *Timelapse microscopy of p0 precursors* in her3:mScarlet-NLS, pax3a:GFP embryos

H2B-haloTag expression: Transgenic embryos obtained from a cross between Tg(*her3:mScarlet-NLS*) and TgBAC(*pax3a:GFP*) transgenic fish were injected at the 1-cell stage with *in vitro*-transcribed H2B-haloTag mRNA. Embryos were dechorionated at 1-2 somite-stage and incubated with JF635 haloTag ligand for 15 min prior to the start of imaging.

Microscopy: Embryos were mounted for dorsal imaging of the neural plate and imaged at 28 C. 3D confocal images were acquired in the H2B-HaloTag channel at 1.5 min intervals to enable cell tracking and GFP/mScarlet channels at 7.5 min intervals to mitigate photo-bleaching of reporter fluorescence.

Analysis: Nuclei were identified and tracked in a semi-automated fashion using the Trackmate 7 plugin within Fiji. Tracks were manually curated within Trackmate prior to downstream analysis. Custom MATLAB scripts were subsequently used to identify GFP+, Scarlet+ cells and analyze cell tracks.

## References

1. Sladitschek, H. L. et al. MorphoSeq: Full single-cell transcriptome dynamics up to gastrulation in a chordate. Cell 181, 922–935.e21 (2020).

2. Guignard, L. et al. Contact area-dependent cell communication and the morphological invariance of ascidian embryogenesis. Science 369, eaar5663 (2020).

3. Xiong, F. et al. Interplay of cell shape and division orientation promotes robust morphogenesis of developing epithelia. Cell 159, 415–427 (2014).

4. Serra, M. et al. A mechanochemical model recapitulates distinct vertebrate gastrulation modes. Sci. Adv. 9, eadh8152 (2023).

5. Fulton, T. et al. Long-term in toto cell tracking using lightsheet microscopy of the zebrafish tailbud. Wellcome Open Res. 3, 163 (2018).

6. McDole, K., et al. In Toto Imaging and Reconstruction of Post-Implantation Mouse Development at the Single-Cell Level. Cell 175, 859-876.e33 (2018).

7. Reversade, B. & De Robertis, E. M. Regulation of ADMP and BMP2/4/7 at opposite embryonic poles generates a self-regulating morphogenetic field. Cell 123, 1147–1160 (2005).

8. Collins, Z., et al. A Scube2-Shh feedback loop links morphogen release and spread to morphogen signaling to enable scale invariant patterning of the ventral neural tube. bioRxiv (2018) doi:10.1101/469239.

9. Zinski, J. et al. Systems biology derived source-sink mechanism of BMP gradient formation. Elife 6, (2017).

10. Greenfeld, H., Lin, J. & Mullins, M. C. The BMP signaling gradient is interpreted through concentration thresholds in dorsal–ventral axial patterning. PLoS Biol. 19, e3001059 (2021).

11. Balaskas, N. et al. Gene regulatory logic for reading the Sonic Hedgehog signaling gradient in the vertebrate neural tube. Cell 148, 273–284 (2012).

12. Dessaud, E. et al. Interpretation of the sonic hedgehog morphogen gradient by a temporal adaptation mechanism. Nature 450, 717–720 (2007).

13. Jaeger, J. et al. Dynamic control of positional information in the early Drosophila embryo. Nature 430, 368–371 (2004).

14. Briscoe, J. & Small, S. Morphogen rules: design principles of gradient-mediated embryo patterning. Development 142, 3996–4009 (2015).

15. Rogers, K. W., ElGamacy, M., Jordan, B. M. & Müller, P. Optogenetic investigation of BMP target gene expression diversity. Elife 9, e58641 (2020).

16. Zagorski, M. et al. Decoding of position in the developing neural tube from antiparallel morphogen gradients. Science 356, 1379–1383 (2017).

17. Gregor, T., Tank, D. W., Wieschaus, E. F. & Bialek, W. Probing the limits to positional information. Cell 130, 153–164 (2007).

18. Xiong, F. et al. Specified neural progenitors sort to form sharp domains after noisy Shh signaling. Cell 153, 550–561 (2013).

19. Wartlick, O. et al. Dynamics of Dpp signaling and proliferation control. Science 331, 1154–1159 (2011).

20. Kicheva, A. et al. Coordination of progenitor specification and growth in mouse and chick spinal cord. Science 345, 1254927 (2014).

21. Lai, H. C., Seal, R. P. & Johnson, J. E. Making sense out of spinal cord somatosensory development. Development 143, 3434–3448 (2016).

22. Cohen, M., Page, K. M., Perez-Carrasco, R., Barnes, C. P. & Briscoe, J. A theoretical framework for the regulation of Shh morphogen-controlled gene expression. Development 141, 3868–3878 (2014).

23. Wine-Lee, L. et al. Signaling through BMP type 1 receptors is required for development of interneuron cell types in the dorsal spinal cord. Development 131, 5393–5403 (2004).

24. Tozer, S., Le Dréau, G., Marti, E. & Briscoe, J. Temporal control of BMP signalling determines neuronal subtype identity in the dorsal neural tube. Development 140, 1467–1474 (2013).

25. Andrews, M. G. et al. BMPs direct sensory interneuron identity in the developing spinal cord using signal-specific not morphogenic activities. Elife 6, (2017).

26. Liem, K. F., Jr, Tremml, G., Roelink, H. & Jessell, T. M. Dorsal differentiation of neural plate cells induced by BMP-mediated signals from epidermal ectoderm. Cell 82, 969–979 (1995).

27. Timmer, J. R., Wang, C. & Niswander, L. BMP signaling patterns the dorsal and intermediate neural tube via regulation of homeobox and helix-loop-helix transcription factors. Development 129, 2459–2472 (2002).

28. Lee, K. J., Dietrich, P. & Jessell, T. M. Genetic ablation reveals that the roof plate is essential for dorsal interneuron specification. Nature 403, 734–740 (2000).

29. Rondon, R. et al. Dual transcriptional activities of PAX3 and PAX7 spatially encode spinal cell fates through distinct gene networks. PLoS Biol. 23, e3003448 (2025).

30. Gribble, S. L., Nikolaus, O. B. & Dorsky, R. I. Regulation and function of Dbx genes in the zebrafish spinal cord. Dev. Dyn. 236, 3472–3483 (2007).

31. Zannino, D. A., Downes, G. B. & Sagerström, C. G. prdm12b specifies the p1 progenitor domain and reveals a role for V1 interneurons in swim movements. Dev. Biol. 390, 247–260 (2014).

32. Le Dréau, G. & Martí, E. Dorsal-ventral patterning of the neural tube: a tale of three signals. Dev. Neurobiol. 72, 1471–1481 (2012).

33. Wijgerde, M., McMahon, J. A., Rule, M. & McMahon, A. P. A direct requirement for Hedgehog signaling for normal specification of all ventral progenitor domains in the presumptive mammalian spinal cord. Genes Dev. 16, 2849–2864 (2002).

34. England, S., Batista, M. F., Mich, J. K., Chen, J. K. & Lewis, K. E. Roles of Hedgehog pathway components and retinoic acid signalling in specifying zebrafish ventral spinal cord neurons. Development 138, 5121–5134 (2011).

35. Litingtung, Y. & Chiang, C. Specification of ventral neuron types is mediated by an antagonistic interaction between Shh and Gli3. Nat. Neurosci. 3, 979–985 (2000).

36. Diez del Corral, R., et al. Opposing FGF and retinoid pathways control ventral neural pattern, neuronal differentiation, and segmentation during body axis extension. Neuron 40, 65–79 (2003).

37. Bang, A. G., Papalopulu, N., Goulding, M. D. & Kintner, C. Expression of Pax-3 in the lateral neural plate is dependent on a Wnt-mediated signal from posterior nonaxial mesoderm. Dev. Biol. 212, 366–380 (1999).

38. Mulligan, K. A. & Cheyette, B. N. R. Wnt signaling in vertebrate neural development and function. J. Neuroimmune Pharmacol. 7, 774–787 (2012).

39. Sasai, N., Kutejova, E. & Briscoe, J. Integration of signals along orthogonal axes of the vertebrate neural tube controls progenitor competence and increases cell diversity. PLoS Biol. 12, e1001907 (2014).

40. Akai, J., Halley, P. A. & Storey, K. G. FGF-dependent Notch signaling maintains the spinal cord stem zone. Genes Dev. 19, 2877–2887 (2005).

41. Henrique, D. et al. cash4, a novel achaete-scute homolog induced by Hensen’s node during generation of the posterior nervous system. Genes Dev. 11, 603–615 (1997).

42. Bae, Y.-K., Shimizu, T. & Hibi, M. Patterning of proneuronal and inter-proneuronal domains by hairy- and enhancer of split-related genes in zebrafish neuroectoderm. Development 132, 1375–1385 (2005).

43. Hans, S. et al. her3, a zebrafish member of the hairy-E(spl) family, is repressed by Notch signalling. Development 131, 2957–2969 (2004).

44. Filippi, A. et al. The basic helix-loop-helix olig3 establishes the neural plate boundary of the trunk and is necessary for development of the dorsal spinal cord. Proc. Natl. Acad. Sci. U. S. A. 102, 4377–4382 (2005).

45. Shkumatava, A., Fischer, S., Müller, F., Strahle, U. & Neumann, C. J. Sonic hedgehog, secreted by amacrine cells, acts as a short-range signal to direct differentiation and lamination in the zebrafish retina. Development 131, 3849–3858 (2004).

46. Wagner, D. E. et al. Single-cell mapping of gene expression landscapes and lineage in the zebrafish embryo. Science 360, 981–987 (2018).

47. Choi, H. M. T. et al. Third-generation in situ hybridization chain reaction: multiplexed, quantitative, sensitive, versatile, robust. Development 145, (2018).

48. Kawanishi, T., Sushida, T., Tsai, T. Y.-C., Takeda, H. & Megason, S. G. Formation control between leader and migratory follower tissues allows coordinated growth. Sci. Adv. 11, eads2310 (2025).

49. Wan, Y. et al. Whole-embryo spatial transcriptomics at subcellular resolution from gastrulation to organogenesis. Science 391, eadt3439 (2026).

50. Cornell, R. A. & Eisen, J. S. Delta signaling mediates segregation of neural crest and spinal sensory neurons from zebrafish lateral neural plate. Development 127, 2873–2882 (2000).

51. Thorpe, C. J., Weidinger, G. & Moon, R. T. Wnt/beta-catenin regulation of the Sp1-related transcription factor sp5l promotes tail development in zebrafish. Development 132, 1763–1772 (2005).

52. Esterberg, R., Delalande, J.-M. & Fritz, A. Tailbud-derived Bmp4 drives proliferation and inhibits maturation of zebrafish chordamesoderm. Dev. Biol. 319, 580 (2008).

53. Takke, C., Dornseifer, P., Weizsäcker, E., v & Campos-Ortega, J. A. her4, a zebrafish homologue of the Drosophila neurogenic gene E(spl), is a target of NOTCH signalling. Development 126, 1811–1821 (1999).

54. Babaryka, A., Kühn, E. & Köster, R. W. In vivo synthesis of meganuclease for generating transgenic zebrafish Danio rerio. J. Fish Biol. 74, 452–457 (2009).

55. Seger, C. et al. Analysis of Pax7 expressing myogenic cells in zebrafish muscle development, injury, and models of disease. Dev. Dyn. 240, 2440–2451 (2011).

56. Satou, C. et al. Transgenic tools to characterize neuronal properties of discrete populations of zebrafish neurons. Development 140, 3927–3931 (2013).

57. Reifers, F. et al. Fgf8 is mutated in zebrafish acerebellar (ace) mutants and is required for maintenance of midbrain-hindbrain boundary development and somitogenesis. Development 125, 2381–2395 (1998).

58. Gouti, M. et al. A gene regulatory network balances neural and mesoderm specification during vertebrate trunk development. Dev. Cell 41, 243–261.e7 (2017).

59. Goto, H., Kimmey, S. C., Row, R. H., Matus, D. Q. & Martin, B. L. FGF and canonical Wnt signaling cooperate to induce paraxial mesoderm from tailbud neuromesodermal progenitors through regulation of a two-step epithelial to mesenchymal transition. Development 144, 1412–1424 (2017).

60. Wilson, V., Olivera-Martinez, I. & Storey, K. G. Stem cells, signals and vertebrate body axis extension. Development 136, 2133–2133 (2009).

61. Gouti, M. et al. In vitro generation of neuromesodermal progenitors reveals distinct roles for wnt signalling in the specification of spinal cord and paraxial mesoderm identity. PLoS Biol. 12, e1001937 (2014).

62. Garriock, R. J. et al. Lineage tracing of neuromesodermal progenitors reveals novel Wnt-dependent roles in trunk progenitor cell maintenance and differentiation. Development 142, 1628–1638 (2015).

63. Fulton, T., Verd, B. & Steventon, B. The unappreciated generative role of cell movements in pattern formation. R. Soc. Open Sci. 9, 211293 (2022).

64. Fulton, T. et al. Cell rearrangement generates pattern emergence as a function of temporal morphogen exposure. bioRxiv (2021) doi:10.1101/2021.02.05.429898.

65. Tsai, T. Y.-C. et al. An adhesion code ensures robust pattern formation during tissue morphogenesis. Science 370, 113–116 (2020).

66. Duval, N., Vaslin, C., Barata, T., Nédélec, S. & Ribes, V. BMP4 patterns Smad activity and generates stereotyped cell fate organisation in spinal organoids. bioRxiv (2019) doi:10.1101/511899.

67. Pierani, A., Brenner-Morton, S., Chiang, C. & Jessell, T. M. A sonic hedgehog-independent, retinoid-activated pathway of neurogenesis in the ventral spinal cord. Cell 97, 903–915 (1999).

68. Benzinger, D. & Briscoe, J. Investigating morphogen and patterning dynamics with optogenetic control of morphogen production. Dev. Cell 60, 3421–3430.e6 (2025).

69. Gard, C. et al. Pax3- and Pax7-mediated Dbx1 regulation orchestrates the patterning of intermediate spinal interneurons. Dev. Biol. 432, 24–33 (2017).

70. Pierani, A. et al. Control of interneuron fate in the developing spinal cord by the progenitor homeodomain protein Dbx1. Neuron 29, 367–384 (2001).

71. Diez del Corral, R., Breitkreuz, D. N. & Storey, K. G. Onset of neuronal differentiation is regulated by paraxial mesoderm and requires attenuation of FGF signalling. Development 129, 1681–1691 (2002).

72. Spiess, K. et al. Approximated gene expression trajectories for gene regulatory network inference on cell tracks. iScience 27, 110840 (2024).

73. Azambuja, A. P., Rothstein, M., Kanno, T. Y. & Simoes-Costa, M. Spatial patterning of the epigenome during vertebrate gastrulation. Nat. Commun. 17, 800 (2025).

74. Lange, M. et al. A multimodal zebrafish developmental atlas reveals the state-transition dynamics of late-vertebrate pluripotent axial progenitors. Cell 187, 6742–6759.e17 (2024).

75. Wan, Y., et al. Whole-embryo Spatial Transcriptomics at Subcellular Resolution from Gastrulation to Organogenesis. bioRxiv (2024) doi:10.1101/2024.08.27.609868.

76. Kim, Y. J., et al. Zebrahub-Multiome: Uncovering Gene Regulatory Network Dynamics During Zebrafish Embryogenesis. bioRxiv (2024) doi:10.1101/2024.10.18.618987.

77. Kinkhabwala, A. et al. A structural and functional ground plan for neurons in the hindbrain of zebrafish. Proc. Natl. Acad. Sci. U. S. A. 108, 1164–1169 (2011).

78. Wolny, A. et al. Accurate and versatile 3D segmentation of plant tissues at cellular resolution. Elife 9, (2020).

79. Ershov, D. et al. TrackMate 7: integrating state-of-the-art segmentation algorithms into tracking pipelines. Nat. Methods 19, 829–832 (2022).

80. Wolf, F. A., Angerer, P. & Theis, F. J. SCANPY: large-scale single-cell gene expression data analysis. Genome Biol. 19, (2018).

81. Shen, W.-K. et al. AnimalTFDB 4.0: a comprehensive animal transcription factor database updated with variation and expression annotations. Nucleic Acids Res. 51, D39–D45 (2023).

82. Schindelin, J. et al. Fiji: an open-source platform for biological-image analysis. Nat. Methods 9, 676–682 (2012).

83. Provost, E. et al. Functional correlates of mutations in beta-catenin exon 3 phosphorylation sites. J. Biol. Chem. 278, 31781–31789 (2003).

