## Supplementary Figures and Legends for "Neural plate morphogenesis and pre-patterning enable specification of intermediate progenitors in the spinal cord"

### Supplementary Figure 1

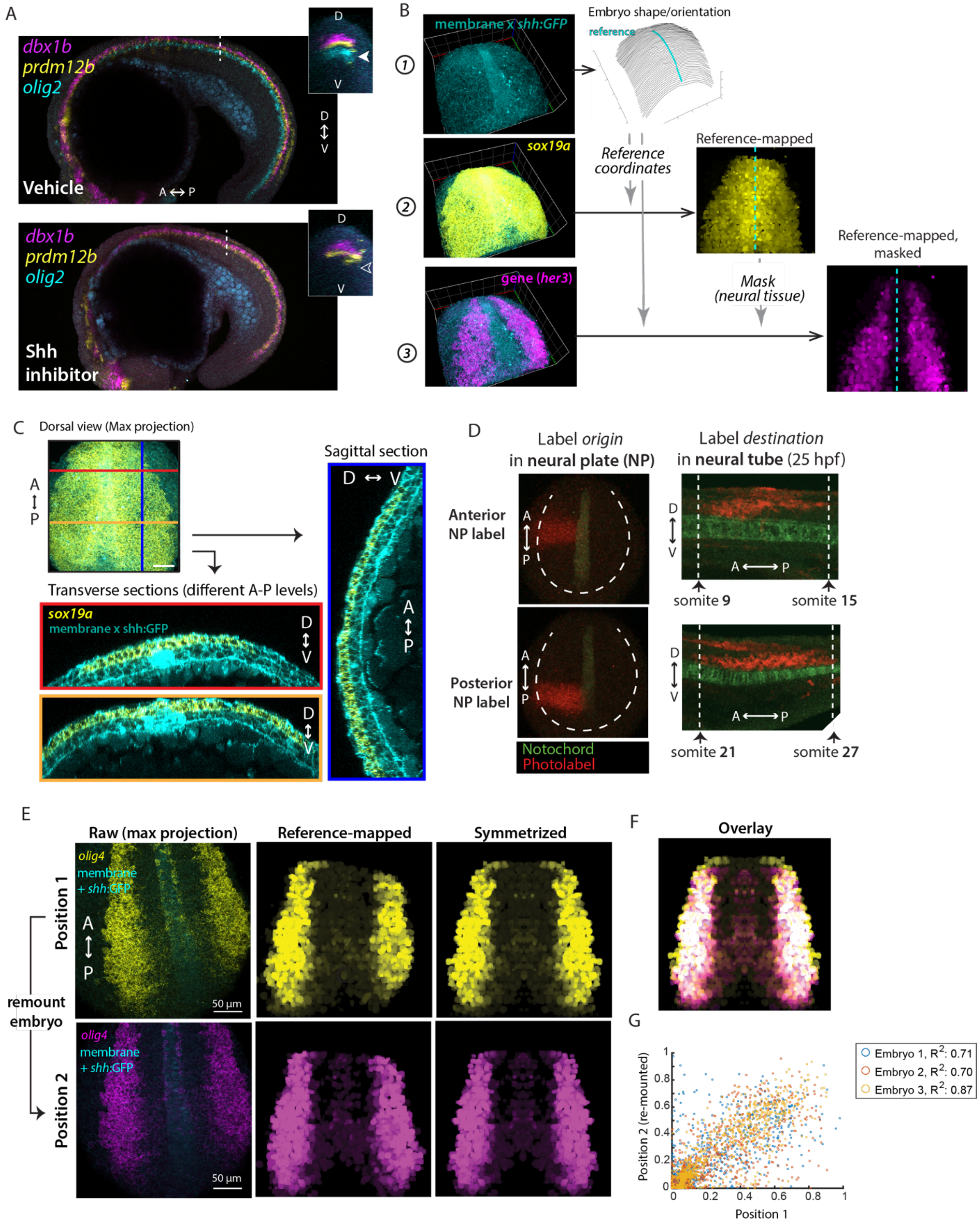


Supplementary Figure 1. **Overview of reference mapping measurements**

(A) Expression of markers for p0 (*dbx1b*, magenta), p1 (*prdm12b*, yellow) and pMN (*olig2*, cyan) neural progenitors at the 18 somite-stage in embryos treated with an inhibitor of Shh signaling (bottom) or a vehicle control (top) starting from 50% epiboly. (Insets) Transverse optical sections at the anterior-posterior position marked by the dotted lines. Filled arrowhead in the top inset points to the ventrally located pMN progenitor domain (cyan) in the vehicle control, while the empty arrowhead in the bottom inset indicates loss of the pMN marker.

(B) Workflow for reference-mapping gene expression in a representative embryo. Measurements are typically made in embryos expressing a membrane-localized mNeongreen fluorescent protein ('mem-Neongreen') and a *shh:GFP* reporter that labels the notochord. (1) (*Left*) 3D-rendering of mem-Neongreen and *shh*:GFP fluorescence signal. (*Right*) mem-Neongreen fluorescence is used to calculate the surface of the embryo (mesh) and to segment individual cells. The midline of the embryos (cyan), annotated based on shh:GFP fluorescence, is used to calculate the orientation of the embryo. (2) (*Left*) 3D-rendering of neural marker *sox19a* HCR RNA-FISH signal (magenta) overlaid on membrane/notochord fluorescence (cyan). (*Right*) Reference-based map of *sox19a*, calculated based on embryo orientation, with midline indicated by dashed line. (3) (*Left*) 3D-rendering of *her3* HCR RNA-FISH signal (magenta) overlaid on membrane/notochord fluorescence (cyan). (*Right*) Reference-based expression of *her3* (magenta), masked using the *sox19a* reference-map in (2).

(C) (*inset*) Maximum projection of 3D image showing membrane-localized mNeongreen and shh:*GFP* reporter fluorescence (cyan) and *sox19a* HCR RNA-FISH (yellow). The image was optically sectioned along the indicated colored lines. (*Right*) Orthogonal planes along lines indicated in inset image. Outline colors of boxes correspond to line colors.

(D) (Left) Dorsal views of neural plate in two 5 somite-stage embryos that constitutively expressed the photo-convertible fluorescent protein kikGR (green), after photo-conversion of indicated regions (red). Dashed line indicates the approximate outlines of the neural plate. (Right) Lateral view showing location of corresponding photo-converted regions at 25 hpf (hours post fertilization). Dashed vertical lines show approximate positions of the indicated somites.

(E) Maximum projection images and reference-mapped versions of *olig4* expression in the same embryo, imaged in two different orientations. The embryo was imaged ('Position 1', yellow), and then re-mounted before imaging ('Position 2', magenta). (Left) Maximum projection images of olig4 (yellow/magenta) and mem-mNeongreen and Shh:GFP fluorescence (cyan). (Middle) Reference-based maps of *olig4* expression. (Right) 'Symmetrized' reference-map obtained by reflecting the side of the neural plate of which more is captured within the field of view.

(F) Overlay of *olig4* reference maps for images acquired in the two orientations in (E).

(G) Scatter plot comparing *olig4* HCR RNA-FISH signal intensity between corresponding pixels of pairs of reference maps generated from imaging the same embryos before and after adjusting their orientation as in (E). ‘Embryo 3’ corresponds to the data shown in (E-F).

### Supplementary Figure 2

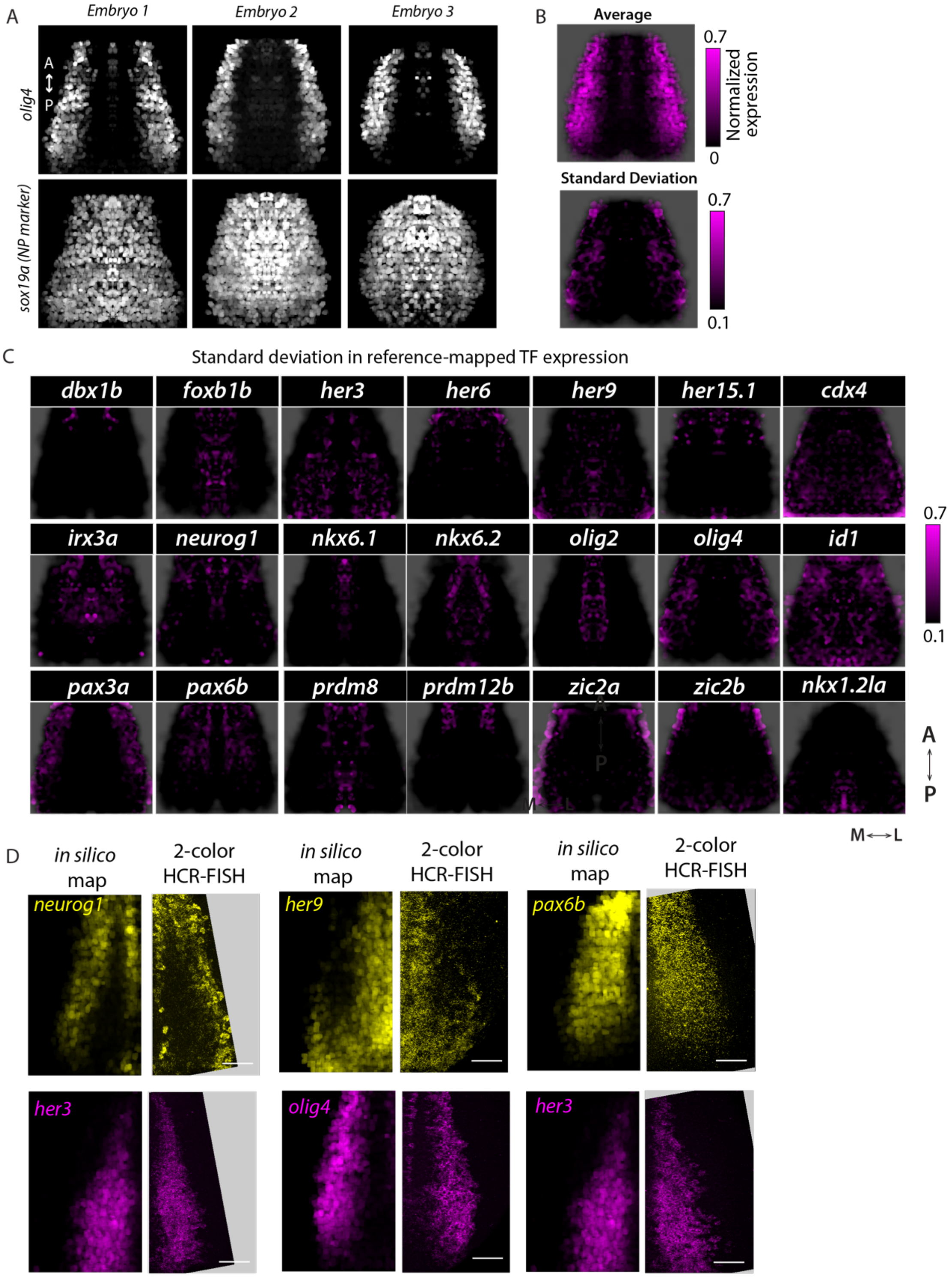


Supplementary Figure 2. **Analysis of embryo-to-embryo variability in reference maps**

(A-B) Calculating average TF expression: an example. (A) Reference-mapped expression of *olig4* (top) and *sox19a* (bottom) in three individual embryos. Signal is normalized to maximum expression in the image. (B) Mean (top) and standard deviation (bottom) of *olig4* reference-maps.
(C) Standard deviation of reference-mapped expression of the indicated transcription factors (related to Figure 1C). The black background corresponds to the neural plate, co-labeled in each measurement using *sox19a*. Each map represents data from n = 3 embryos, each symmetrized and normalized to the maximal detected signal within the image.
(D) Individual channels from Figure 1D, comparing average reference-mapped expression (left panels) with co-expression in representative embryos detected using HCR RNA-FISH. Scale bar indicates 50 μm.

**Supplementary Figure 3**

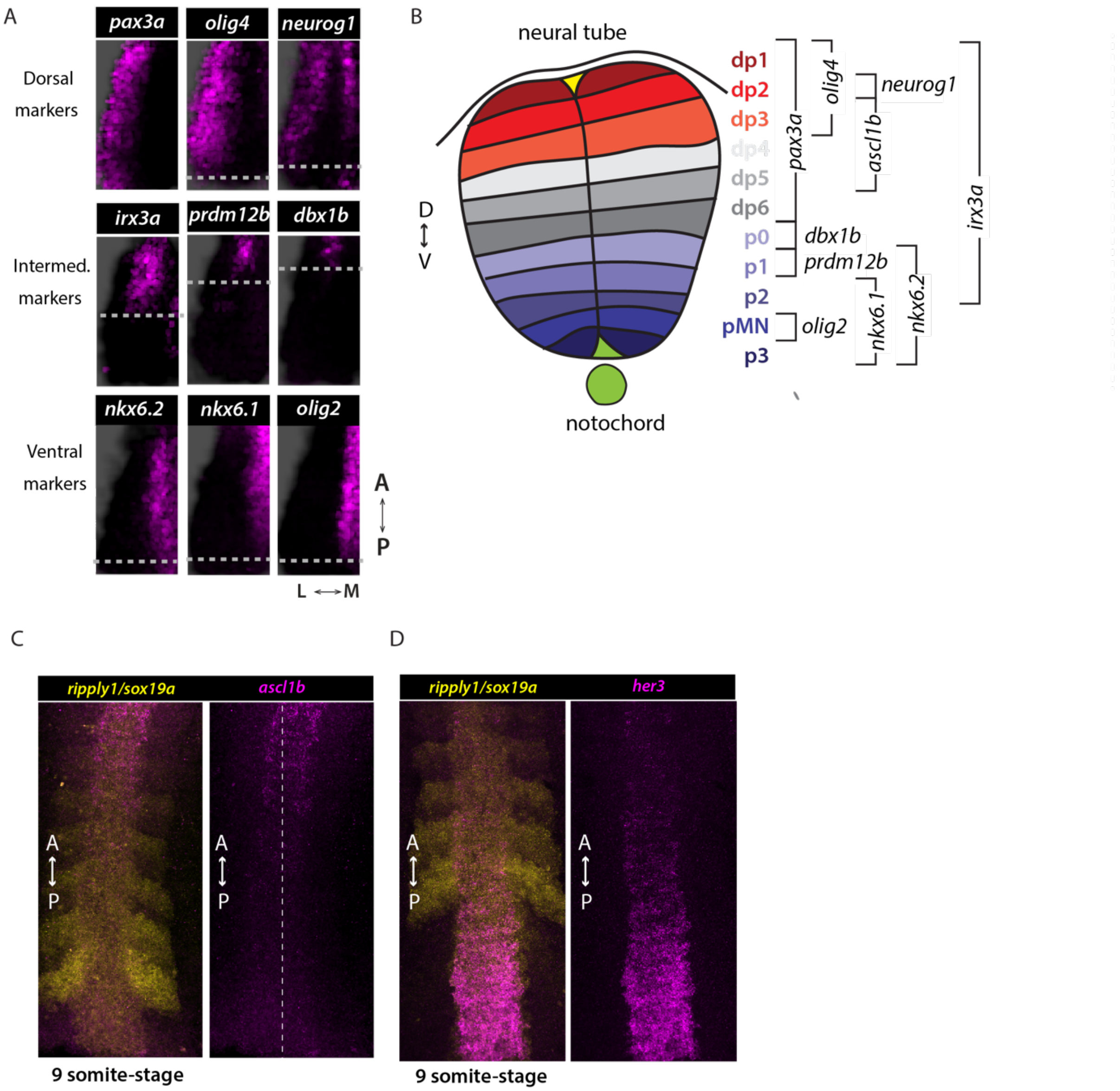


Supplementary Figure 3. **Dorsal-ventral markers in the neural tube are expressed at different antero-posterior levels in the neural plate**

(A) Comparison of the posterior extent of TFs that mark different dorsal-ventral domains within the future neural tube (NT) (see B). Dashed line indicates approximate posterior boundary in average reference-mapped expression. No line is shown in cases in which expression can be observed in the posterior-most extent of the map. Same data as Figure 1D. (B) Schematic indicating expression of indicated TFs across different neural progenitor domains in the neural tube.

(C) Representative maximum projection image showing that expression of the dorsal neural tube marker *ascl1b* (magenta) is only initiated anterior to somite formation (*ripply1* expression, yellow).
(D) Representative maximum projection image showing that *her3* is expressed posterior to the somite formation (*ripply1* expression, yellow) but is extinguished at more anterior positions.

### Supplementary Figure 4

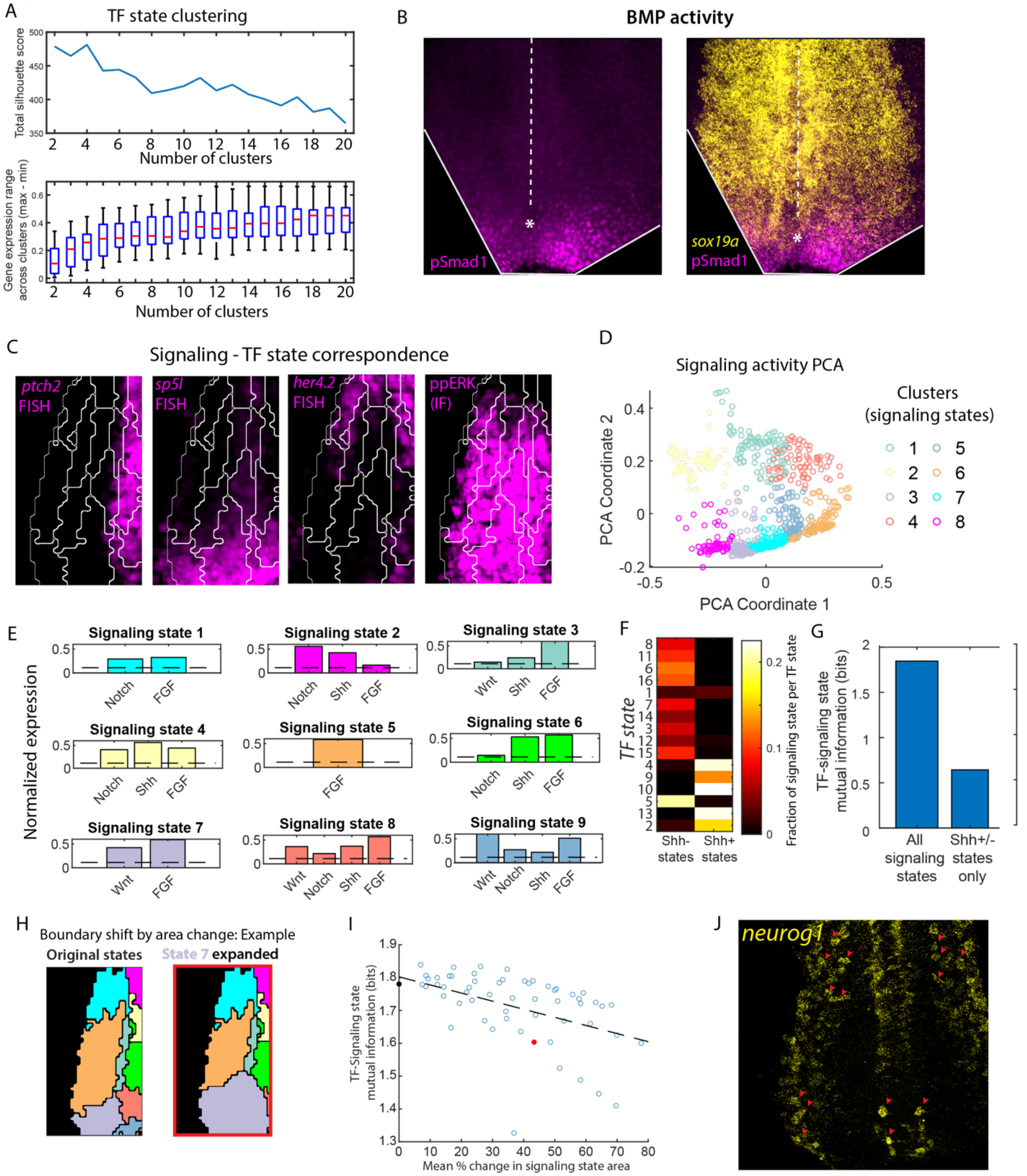


Supplementary Figure 4. **Supporting data for identification of TF and signaling states in the neural plate**

(A) (Top) Silhouette score vs. number of clusters for the k-means clustering analysis in Figure 2. (Bottom) Boxplots showing the range of expression for TFs across clusters. For each TF, this is calculated by averaging its expression within each cluster and then calculating the difference between the maximum and minimum values of this value. Higher values indicate that TF expression is enriched (or depleted) within some clusters and suggests better segregation of superpixels expressing different levels of that TF. Note that decrease in silhouette score is compensated by an increase in this separation ability until ~16 clusters, which is the value chosen for *k*-means clustering in Figure 2.

(B) Representative example showing immunofluorescence for phosphorylated-Smad1 ('pSmad1', magenta) and HCR RNA-FISH for the neural marker *sox19a* (yellow) in the neural plate region of the embryo. Maximum projection of a 3D image is shown. Dashed line indicates approximate midline of the neural plate, based on a co-expressed notochord marker (not shown), while the asterisk marks its posterior end. Note exclusion of pSmad1 from the neural plate.

(C) Outlines of TF states from Figure 2C overlaid on reference maps of Shh, Wnt, Notch and FGF activity (same as Figure 2E).

(D) Scatter plot showing principal component values for signaling activity levels in superpixels, colored by the clusters they are assigned to after *k*-means clustering.

(E) Mean levels of signaling targets within the individual states shown in Figure 2E. Only pathways with mean target levels above a threshold (dashed line) are shown.

(F) Heatmap showing how signaling states characterized by high Shh activity ('Shh+') or low activity ('Shh-') are distributed across TF states. Each column shows the fractional distribution of the corresponding signaling state across all the TF states.

(G) Barplots comparing mutual information between all signaling states and TF states vs. states distinguished only by Shh activity levels ('Shh+/- states') and TF states.

(H) (Right) Example of modified signaling state map generated by altering the area of one state (State 7, purple). (Left) Original signaling state map.

(I) Scatter plot showing relationship between TF-signaling state mutual information and the extent to which the signaling state map is modified, measured by the mean percentage change in the area of signaling states. The dashed line represents the linear fit to the data. The black point corresponds to the original signaling state map, while the red point corresponds to the example shown in (H).

(J) Maximum intensity projection of *neurog1* expression in the neural plate, showing salt-and-pepper distribution. Red arrowheads highlight some isolated *neurog1*+ cells surrounded by cells expressing low or no *neurog1*.

### Supplementary Figure 5

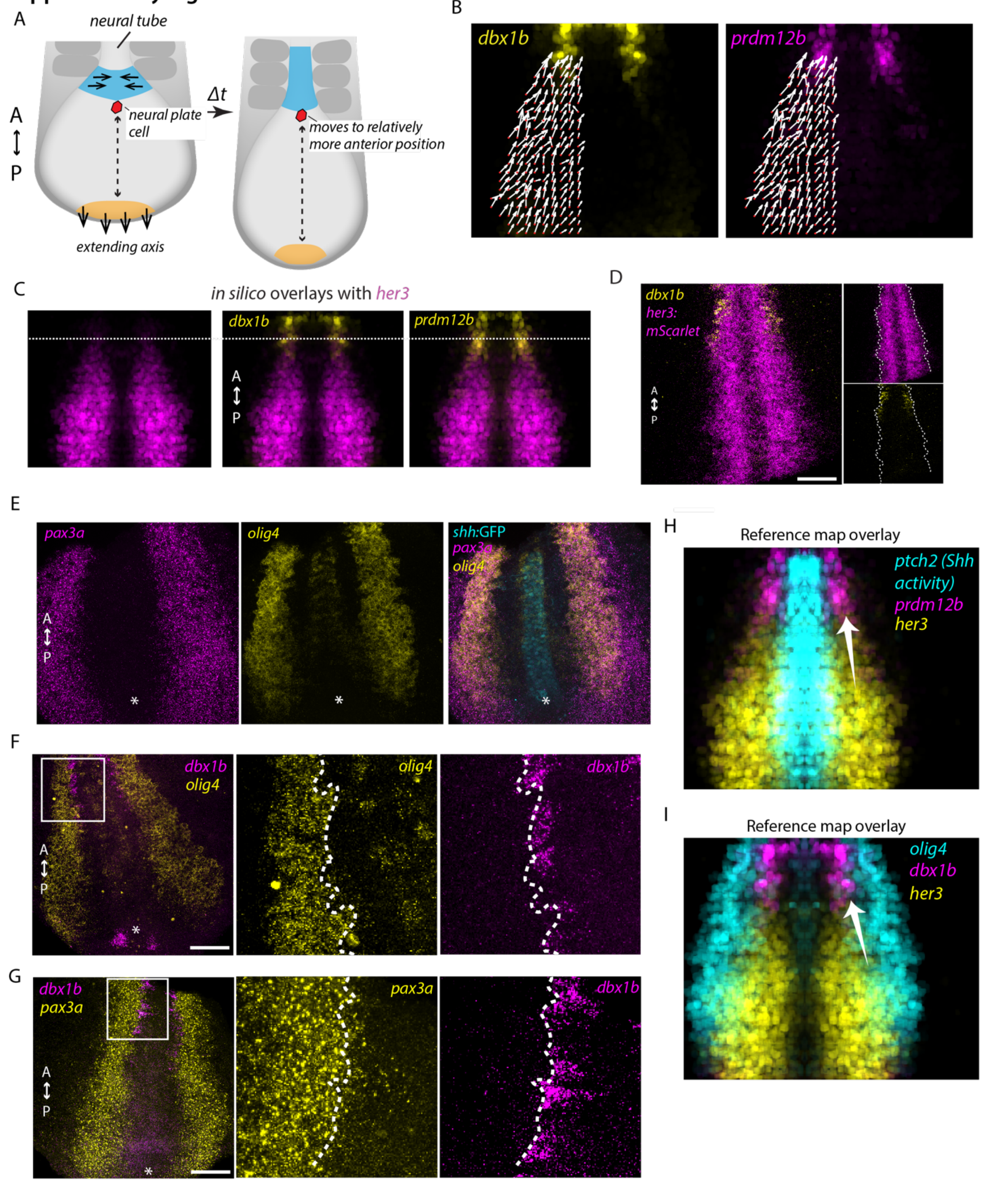


Supplementary Figure 5. ***p0 and p1 cells originate from specific precursor states characterize by* olig4, her3, pax3a+ *expression***

(A) (Schematic) During axis elongation, the neural plate converges in more anterior regions (blue) forming the neural tube, while extending in posterior regions (yellow). Due to this relative difference in the dominant directions of cell movement along the antero-posterior axis, most cells in the neural plate shift to more anterior positions relative to the posterior end as time progresses.

(B) Overlay of reference map of cell motion (restricted to *her3* expression domain, same as Figure 3D) with maps of *dbx1b* (left) or *prdm12b* (right) expression.

(C) Reference map of *her3* expression (magenta) or *in silico* overlays of *her3* with reference maps of *dbx1b* (middle) or *prdm12b* (right); same data as Figure 1D. Dashed line indicates approximate anterior limit of *her3* expression.

(D) Overlay of *dbx1b* expression (yellow) and *her3*:*mScarlet-NLS* reporter expression (magenta), measured by HCR RNA-FISH. Images represent maximum intensity projections of 3D images. Dashed white lines indicate approximate lateral boundaries of the *her3*:*mScarlet-NLS* expression. Note the *dbx1b* expression is contained within the *her3*:*mScarlet-NLS* expression, and located at the lateral boundary. Scale bar represents 50 $\mu$m.

(E) Representative example of *pax3a* (magenta) and *olig4* (yellow) expression, measured using HCR RNA-FISH, in a single transgenic embryo expressing a *shh:GFP* reporter, which marks the notochord. Maximum projection of 3D image is shown. The asterisk marks the posterior end of of the notochord. Scale bars represent 50 $\mu$m.

(F) (*Left*) Representative example of *dbx1b* (magenta) and *olig4* (expression) expression, measured using HCR RNA-FISH, in a single embryo. Maximum projection of 3D image is shown. White box indicates a region of *dbx1b* expression, magnified on the right. (*Middle, Right) dbx1b* (magenta) and *olig4* (yellow) expression in the region outlined by the white box on the left. The dashed lines indicate the medial boundary of *olig4* expression. Note that *dbx1b* is only expressed outside the *olig4* boundary. Scale bar represents 50 $\mu$m.

(G) (*Left*) Representative example of *dbx1b* (magenta) and *pax3a* (expression) expression, measured using HCR RNA-FISH, in a single embryo. Maximum projection of 3D image is shown. White box indicates a region of *dbx1b* expression, magnified on the right. (*Middle, Right) dbx1b* (magenta) and *pax3a* (yellow) expression in the region outlined by the white box on the left. The dashed lines indicate the medial boundary of *olig4* expression. Note that *dbx1b* is only expressed outside the *pax3a* boundary. Scale bar represents 50 $\mu$m.

(H, I) *in silico* overlays of reference maps for indicated genes indicating proposed origin for (H) p1 (*prdm12b*+) cells from her3 cells lateral to the Shh activity region (marked by *ptch2l* expression) and (I) p0 (*dbx1b*+) cells from *olig4/her3* -coexpressing cells at the lateral boundary of *her3* expression.

### Supplementary Figure 6

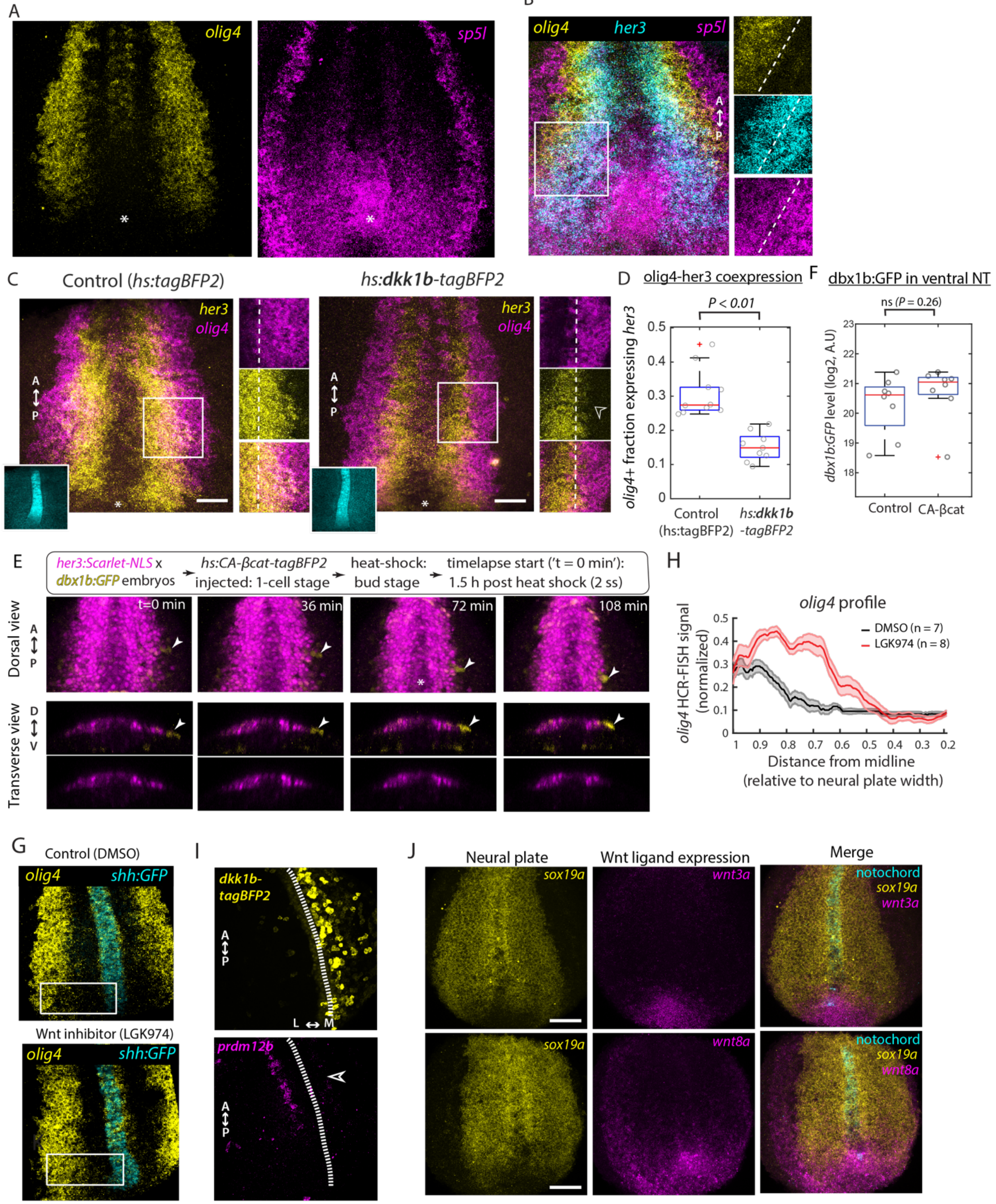


**Supplementary Figure 6. Tailbud-derived Wnt signaling defines p0 and p1**

(A) Representative example of *sp5l* (magenta) and *olig4* (yellow) expression, measured using HCR RNA-FISH, within an individual embryo (same as Figure 4B). Maximum projection of 3D image is shown.

(B) (*Left*) Representative example showing expression of *olig4, her3* and *sp5l*, measured using HCR RNA-FISH, in the neural plate region of an embryo. Maximum projection of a 3D image is shown. White box indicates a region of *olig4, her3* and *sp5l* coexpression, magnified on the right. (*Middle, Right) dbx1b* (magenta) and *olig4* (yellow) expression in the region outlined by the white box on the left. The dashed lines indicate the medial boundary of *olig4* expression. Note that *dbx1b* is only expressed outside the *olig4* boundary. Scale bar represents 25 um.

(C-D) Effect of ectopic expression of *dkk1b* (Wnt signaling inhibitor) on *olig4/her3* co-expression. (C) (*Left*) Representative examples showing expression of *olig4* (magenta) and *her3* (yellow), measured using HCR RNA-FISH, in the neural plate region of transgenic *shh:GFP, mem-mNeongreen* embryos expressing heat-shock inducible ('hs') *dkk1b-tagBFP2* (bottom) or *tagBFP2* control (top). Maximum projection of 3D images is shown. Asterisks indicate approximate location of the posterior end of the notochord, based on the fluorescent marker *shh:GFP* (cyan, insets). White boxes indicate regions where *olig4* and *her3* are normally co-expressed (see Figure 4B), magnified in the panels on the right. Scale bars represent 25 um. (*Right panels*) *olig4* (magenta) and *her3* (yellow) expression in the regions outlined by the white boxes in the corresponding images on the left. The dashed lines indicate the medial boundary of *olig4* expression. Note expression of *her3* within the *olig4*+ region in the control condition (*top*) but not in the *dkk1b* condition (*bottom*).

(D) Boxplots showing the fraction of *olig4*+ cells co-expressing *her3* in transgenic *shh:GFP, mem-mNeongreen* embryos after ectopic expression of *dkk1b-tagBFP2* (n=9 embryos) or *tagBFP2* control (n=10 embryos). *olig4* and *her3* expression were measured using HCR RNA-FISH and membrane mNeongreen fluorescence was used to calculate co-expression in individual cells. *P*-values calculated using Student's t-test.

(E) (*Top*) Experiment scheme for timelapse imaging of ectopic induction of dbx1b-GFP cells following *CA-βcat-tagBFP2* expression. (*Bottom*) Timecourse showing ectopic induction of a *dbx1b:GFP*+ cell (arrowhead, reporter fluorescence in yellow) from a region lateral to the expression domain of *her3:mScarlet-NLS* (magenta) following induction of *CA-βcat-tagBFP2*. Top panels represent 3D-rendered images from a dorsal perspective, while bottom panels show transverse cross-sections at the axial level of the induced cell.

(F) Boxplots showing total GFP fluorescence <u>within</u> the mScarlet-NLS+ region, in the neural tube of transgenic *dbx1b:GFP, her3:mScarlet-NLS* embryos, either ectopically expressing *CA-βcat-tagBFP2* (n=8 embryos) or *tagBFP2* control (n=8 embryos). Same experiment as Figure 5H, I. *P*-value calculated by Student's t-test.

(G-H) Effect of Wnt signaling inhibition on *olig4* expression. (G) Representative images showing expression of *olig4* (yellow) in embryos treated either with a control (top) or with the Wnt signaling inhibitor LGK974 (bottom). Embryos also express the notochord marker *shh:*GFP (cyan). This marker is used to automatically choose a region at the posterior end of the neural plate (white box) where the *olig4* medio-lateral expression profile is calculated for plotting in (H).

(H) Median medio-lateral expression profile of *olig4* in the posterior neural plate, after 3h treatment with the Wnt signaling inhibitor LGK974 (red) or a vehicle control DMSO (black). Shaded areas indicate S.E.M.

(I) Representative example showing expression of *prdm12b* (magenta) in the neural plate in embryos ectopically expressing *dkk1b-tagBFP2* (yellow). Expression was measured using HCR RNA-FISH. Dashed line indicates approximate location of the midline. Empty arrowhead highlights reduction in *prdm12b* expression on the side of the embryo expressing higher levels of *dkk1b-tagBFP2*.

(J) Representative examples showing expression of the Wnt ligands *wnt3a* (top) and *wnt8a* (bottom), with the neural marker *sox19a,* measured by HCR RNA-FISH. Embryos also expressed the notochord marker *shh:*GFP (cyan). Maximum projection of a 3D image is shown. Note expression of Wnt ligands is concentrated posterior to the neural plate and notochord, corresponding to the tailbud of the embryo. Scale bar represents 50 um.

### Supplementary Figure 7

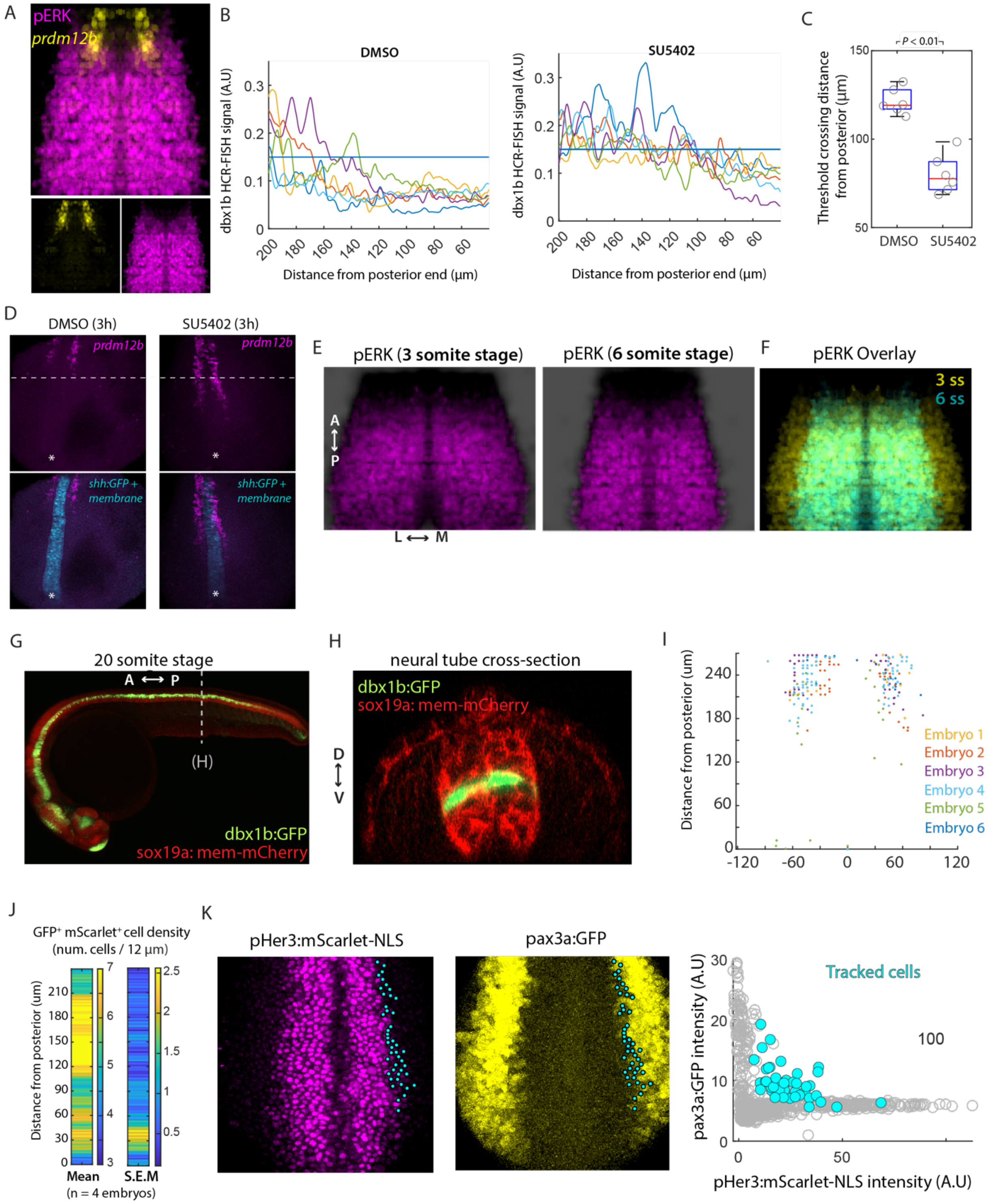


Supplementary Figure 7: **Supporting data for role of convergent extension movements in specifying p0 and p1 neural progenitors**

(A) Overlay of reference maps for *prdm12b* (yellow) and phosphorylated ERK (magenta) from Figure 1D and Figure 5A, respectively.
(B) Axial profiles of *dbx1b* HCR RNA-FISH signal intensity in individual embryos treated with SU5402 or DMSO control, used for calculation of average profiles in Figure 5C. Horizontal line indicates an intensity threshold used for quantification in (C).
(C) Boxplots showing position of threshold crossing in profiles in (B). *P*-value calculated using Student's t-test.
(D) Representative images showing *prdm12b* expression (magenta), measured using HCR RNA-FISH, in transgenic embryos expressing *shh:GFP* and membrane-localized mNeongreen (cyan) after treatment with SU5402 or control (DMSO) for 3 hours. Shown are 3D rendered images from a dorsal perspective. The dashed line indicates the posterior extent of prdm12b in the DMSO-treated embryo, while the asterisk indicates the posterior end of the notochord based on *Shh:GFP* expression.
(E-F) Comparison of neural plate FGF activity at 3 and 6 somite stages ('ss'). (D) Reference maps of phosphorylated ERK (pERK) immunofluorescence (magenta) at 3 somite stage ('3 ss') and 6 somite stage ('6 ss'). Dark background in each image indicates outline of neural plate, based on *sox19a* expression (not shown). Note decrease in medial width of the neural plate between 3 ss and 6 ss due to convergence-extension morphogenesis. Each map represents the average of reference maps from three embryos. (E) Overlay of reference maps shown in (D). Note similar anterior boundary in pERK levels.
(G, H) dbx1b:GFP fluorescence (green) in a representative embryo that co-expresses a sox19a:mem-mCherry neural marker (red) at the 20 somite-stage. (G) Lateral view, anterior to the left. (H) Transverse optical section at the dashed line indicated in (G). Note the narrow one-cell wide extent of the p0 (dbx1b:GFP+) domain.
(I) Reference map positions of *dbx1b*+ cells in individual embryos, used for quantification in Figure 5E.
(J) Heatmaps showing mean and S.E.M of the density of $GFP^{+}mScarlet\text{-}NLS^{+}$ cells in pax3a:GFP, her3:mScarlet-NLS double transgenic embryos cells at different axial levels in the neural plate.
(K) Identification of cells co-expressing GFP and mScarlet-NLS (cyan), tracked in the timelapse analysis of Figure 6H, based on levels of pax3a:GFP (yellow) and her3:Scarlet-NLS (magenta) fluorescence levels.
